# Latrophilin-2 Orchestrates Endothelial Flow-Mediated Smad1/5 Activation via Endoglin and Shank3

**DOI:** 10.1101/2025.06.30.662394

**Authors:** Keiichiro Tanaka, Manasa Chanduri, Hyojin Park, Minghao Chen, Andrew Prendergast, Divyesh Joshi, Yoo-Min Koh, Jay Solanki, Anne Eichmann, Martin A. Schwartz

## Abstract

Fluid shear stress (FSS) from blood flow critically determines vascular morphology and function. FSS activates the Smad1/5 pathway maximally at physiological FSS through the BMP9/10 receptors Alk1 and Endoglin to promote vascular stability. We now report that the adhesion G protein-coupled receptor Latrophilin-2 (Lphn2), which mediates flow activation of the canonical junctional complex, is required for flow-mediated Smad1/5 activation in endothelial cells. Lphn2 physically associates with Endoglin and is required for Smad1/5 activation by FSS but not soluble BMP9. This regulation is independent of G-proteins but requires Lphn2’s C-terminal PDZ binding motif. Endothelial-specific Lphn2 knockout reduces Smad1/5 activation in mice. In zebrafish embryos, loss of flow or Lphn2 knockout similarly reduced endothelial Smad1/5 activation, with no additive effect, indicating that Lphn2 mediates flow-induced activation *in vivo*. APEX2 proximity labeling revealed that the PDZ domain-containing scaffold protein Shank3 associated with Lphn2 under flow. Shank3 knockdown blocked flow-induced Smad1/5 activation *in vitro* and Shank3 global knockout mice exhibited reduced endothelial nuclear Smad1/5 and increased susceptibility to arteriovenous malformations. Lphn2 thus mediates flow-induced Smad1/5 activation and vessel stabilization through the scaffold protein Shank3, independently of G proteins. Together with the accompanying paper on Notch1, we conclude that Lphn2 confers FSS sensitivity to three of the major FSS pathways via distinct mechanisms.

## Introduction

Fluid shear stress (FSS) from blood is sensed by vascular endothelial cells (ECs) to activate numerous signaling and gene expression pathways that govern embryonic vessel development and adult physiology and pathology. In adults, laminar FSS within the physiological range (i.e., near the set point) stabilizes blood vessels whereas FSS above or below the set point induces, respectively, outward or inward remodeling to restore FSS to the desired range. These processes constitute an essential homeostatic mechanism that determines vessel diameter to optimize delivery of oxygen and nutrients to tissues^1^.

EC flow responses are determined by an integrated network of FSS mechanotransduction pathways. One of these involves a mechanosensory complex at cell-cell junctions whose components identified to date include PECAM1, VEGFR2 and 3, VE-cadherin, neuropilin-1 and PlexinD1^2–5^. This complex plays a crucial role in flow-dependent vessel remodeling as well as atherosclerosis susceptibility at regions of disturbed FSS. Recently, latrophilin2 (Lphn2/ADGRL2) was identified as a junctional protein that physically associates with PECAM1 and is required for flow-induced activation of the complex^6^.

A second mechanism for FSS regulation of vascular morphology and function involves bone morphogenetic proteins (BMP) 9 and 10, which signal through the activin-like kinase (ALK)1-Endoglin (Eng) receptor complex to activate the Smad1/5/8 family of transcription factors, with Smad1/5 dominant in ECs^7^. FSS amplifies the activation of Smad1 and 5 by circulating BMPs, coincident with increased association of the type 1 BMP receptor Alk1 and the type 3 receptor Endoglin (Eng)^8^. Active Smad1/5 promote vascular quiescence and stability, with key downstream target genes that induce cell cycle arrest and smooth muscle cell recruitment while inhibiting the VEGFR2-PI3K pathway required for remodeling^8,9^. Smad1/5 activity at later times is maximal at physiological FSS, as high FSS induces expression of BMPER, which suppresses Smad1/5. This reduction in Smad1/5 activity then permits activation of VEGFR2 and PI3K, which are required for homeostatic outward artery remodeling under high FSS^9^. As predicted from this pathway design, mutation of Alk1, Eng or Smad4 leads to vascular malformations in hereditary hemorrhagic telangiectasia (HHT)^8,10^. These lesions are PI3K-dependent and exhibit characteristics of high FSS-induced outward vessel remodeling^11^.

The adhesion G protein-coupled receptor (aGPCR) Lphn2 has recently emerged as a multifunctional signaling molecule beyond its established role in neuronal systems^12^. Lphns contain a large, multidomain extracellular region linked to a seven-transmembrane domain via a GPCR autoproteolysis-inducing (GAIN) domain. Initially identified in neuronal synapse formation and neurotransmitter release, Lphns have broader functions in cell adhesion, signaling, and development^13–15^. The Lphn cytoplasmic tail contains a PDZ-binding motif, enabling interactions with intracellular scaffold proteins and positioning Lphns as potential organizers of signaling complexes^16^. We recently identified Lphn2 as a critical shear stress sensor upstream of PECAM1 at endothelial junctions^6^. FSS activates Gαi2 and Gαq/11 through Lphn2 to initiate the junctional mechanotransduction cascade, establishing Lphn2 as the previously unknown upstream mechanosensor in this pathway. Loss of Lphn2 in zebrafish and endothelial cell knockout (ECKO) in mice led to defects in flow-dependent developmental angiogenesis and arterial morphogenesis and remodeling. Additionally, human mutations in the Lphn2 gene are linked to vascular disease, underscoring relevance to vascular pathophysiology.

In the accompanying study, we found that Lphn2 ECKO increased atherosclerosis in hyperlipidemic mice, opposite to the effect of PECAM1 deletion. This prompted us to investigate whether Lphn2 contributes to additional endothelial FSS-sensing pathways. Here, we report a role for Lphn2 in flow-induced activation of Smad1/5, involving a physical association with Endoglin and interaction the PDZ domain-containing scaffold protein Shank3. Shank3-null mice show increased malformations when challenged by injection of BMP9/10 antibodies. Together, these findings uncover an unexpected G protein-independent signaling function for Lphn2 requiring Shank3 in flow-dependent Smad signaling, vascular biology and disease.

## Results

### Flow-mediated Smad1/5 activation requires Lphn2

Previous work showed that fluid shear stress (FSS) activates the BMP9/10-ALK1-Eng signaling axis, promoting phosphorylation and nuclear translocation of Smad1/5 to support vessel stabilization and endothelial quiescence^8^. ALK1 was found to localize to cell-cell contacts via VE-cadherin, which promotes ALK1-containing receptor complex assembly and downstream signaling in TGF-β-stimulated endothelial cells^17^. This result prompted us to investigate the role of cell-cell adhesion in FSS-induced Alk1-mediated signaling. ECs at varying densities were subjected to laminar shear stress at 12 dynes/cm^2^, the midpoint of the physiological range in HUVECs. Phosphorylation of Smad1/5 was markedly stronger in confluent monolayers than in sparse cells (Figure 1A-B). The effect of cell density persisted in the presence of a potent inhibitor of cell cycle-dependent kinases (Supplemental Figure 1), excluding increased proliferation of sparse cells as the underlying cause. Cell-cell junctions are thus essential for flow-enhanced Smad1/5 signaling. This pathway, however, is reportedly independent of the PECAM1/VE-cadherin/VEGFR2 junctional mechanosensory complex^8^.

**Figure 1:**
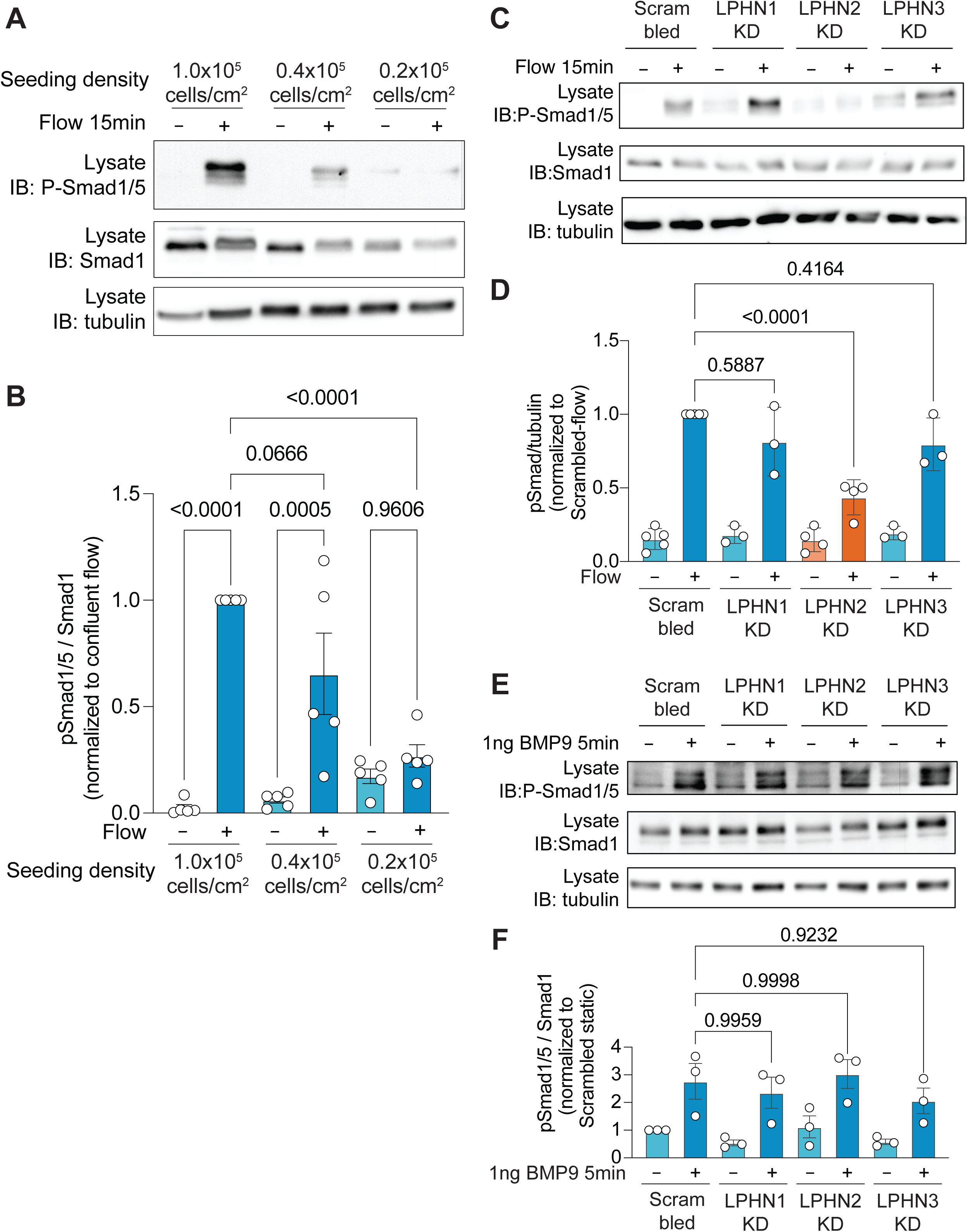
Latrophilin-2 specifically regulates flow-induced Smad1/5 activation. **A)** Density-dependent Smad1/5 phosphorylation in HUVECs. Confluent (1x10^5^ cells/cm^2^), 50% confluent (0.4 x10^5^ cells/cm^2^), and 25% confluent (0.2 x10^5^ cells/cm^2^), without or with 15 minutes laminar flow at 12 dynes/cm^2^. **B**) Quantification of phosphorylated Smad1/5 normalized to total Smad1 in (**A**). Values are means ± S.E.M from n = 5 independent experiments. **C**) Smad1/5 phosphorylation in cells transfected with scrambled siRNA (control) or siRNA targeting Lphn1, Lphn2, or Lphn3, without or after 15 minutes laminar flow. **D**) Quantification of phosphorylated Smad1/5 normalized to tubulin in (**C**). Values are means ± S.E.M from n = 5 independent experiments. **E**) Smad1/5 phosphorylation in control or Lphn1, Lphn2, or Lphn3 knockdown HUVECs treated with or without high dose of BMP9 (1 ng/mL) for 5 minutes. **F**) Quantification of phosphorylated Smad1/5 normalized to total Smad1 in (**E**). Values are means ± S.E.M from n = 3 independent experiments. Statistical significance was assessed using one-way ANOVA with Tukey post hoc tests.

We next addressed the requirement for Latrophilin-2 (Lphn2) as a key component of the junctional flow-sensing machinery. Knockdown of Lphn2, but not its paralogs Lphn1 or Lphn3 (which are minimally expressed in ECs^6^), impaired Smad1/5 activation by FSS in HUVECs (Figure 1C-D). Similar effects were observed in HAECs (Supplementary Figure 2A-C). We also observed reduced flow-induced expression of canonical endothelial Smad1/5 target genes, including *Smad6*, *Id1* and *Jag1*^18^ (Supplementary Figure 2D). By contrast, the response to high dose BMP9 without flow was unaffected (Figure 1E-F). Lphn2 is thus selectively required for flow-but not ligand-induced Smad1/5 activation.

### Flow-mediated Smad 1/5 activation is independent of G-proteins

FSS rapidly activates G-proteins in ECs, with Gαq/11 and Gi proteins playing key roles^6,19–23^. To assess their role in Smad1/5 activation by flow, we selectively depleted Gi and Gq/11 in HUVECs (Figure 2A) and assessed Smad1/5 phosphorylation in response to FSS. To our surprise, knockdown of Gi, Gq/11, or all combined had no effect on flow-induced Smad1/5 phosphorylation (Figure 2B-C).

**Figure 2:**
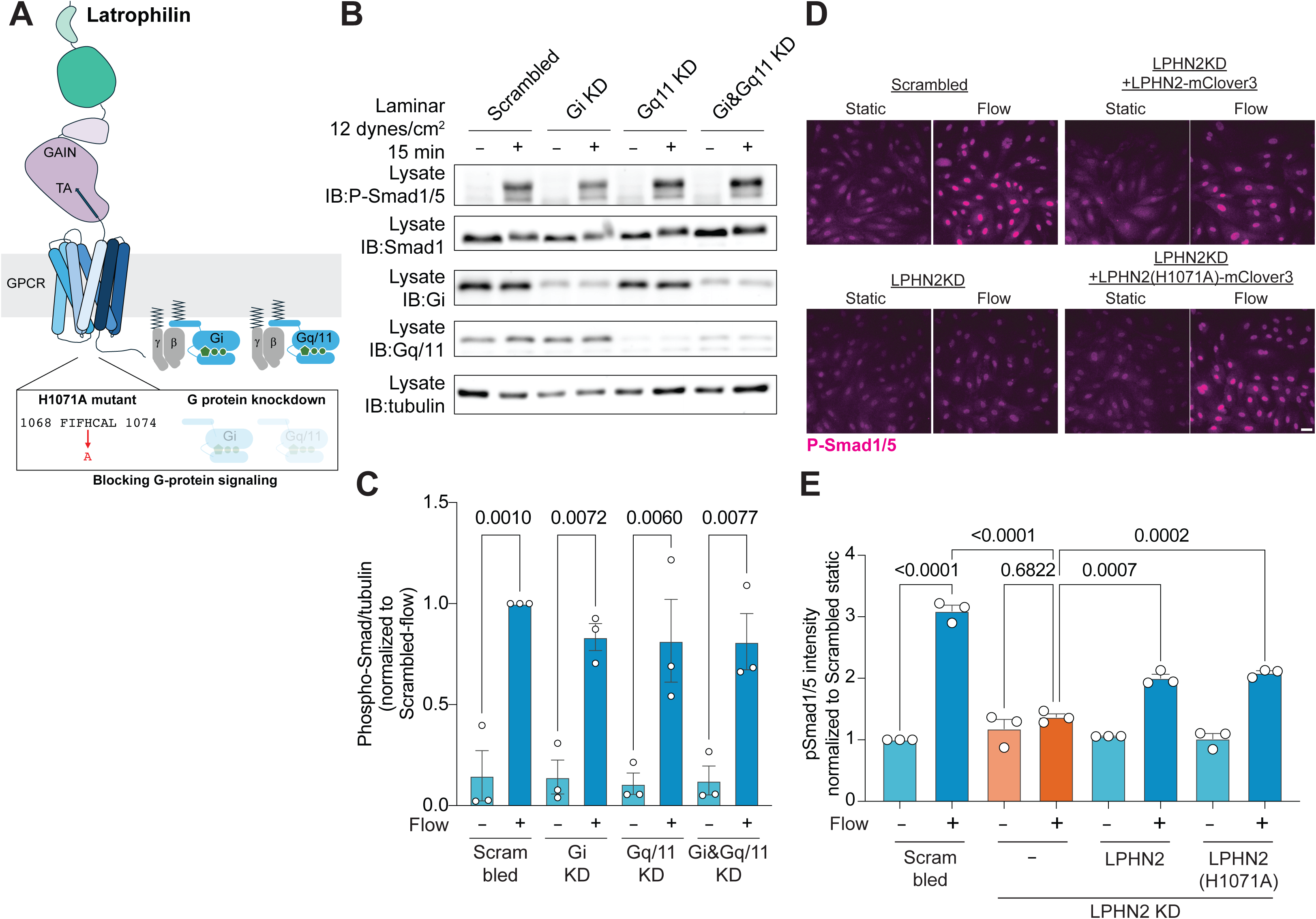
Smad1/5 activation is G protein independent. **A**) Schematic of G proteins and Latrophilin-2 (LPHN2) structure with mutations indicated. **B**) Smad1/5 phosphorylation in HUVECs transfected with scrambled siRNA (control) or siRNA targeting Gαi, Gαq/11, or both, with or without 15 minutes laminar FSS at 12 dynes/cm^2^. **C**) Quantification of phosphorylated pSmad1/5 signal in (**B**) normalized to tubulin. Values are means ± S.E.M. from n = 3 independent experiments. **D**) Nuclear translocation of phosphorylated Smad1/5 in scrambled siRNA (control) and Lphn2 knockdown HUVECs rescued with either wild-type Lphn2 or the G-protein activation-deficient mutant (H1071A), with or without laminar FSS for 30 min. Scale bar, 50 µm. **E**) Quantification of nuclear pSmad1/5 intensity in (**D**). Values are normalized to scrambled static and means ± S.E.M. from three independent experiments. Each experiment quantified 400-800 cells per condition. Statistics were assessed using one-way ANOVA with Tukey post hoc tests.

β-arrestins, which are key mediators of GPCR desensitization pathways, can function as scaffolds for G protein–independent signaling pathways. However, siRNA-mediated depletion of β-arrestin 1 or 2 or both did not alter Smad1/5 activation in response to FSS (Supplementary Figure 3A-C).

To further test G protein-independence, we knocked down endogenous Lphn2 in HUVECs and rescued these cells with either wild-type Lphn2 or the G protein activation-deficient H1071A mutant (Figure 2A)^24^. Re-expression of wild-type and H1071A Lphn2 restored flow-induced activation of Smad1/5 equally well (Figure 2D-E). Together, these results demonstrate that Lphn2’s GPCR activity is dispensable for Smad1/5 activation by flow.

### Lphn2 regulates Smad1/5 activation *in vivo*

To investigate the role of Lphn2 in Smad1/5 activation *in vivo*, we first assessed Smad1/5 activation in ECs of aortas from wild-type (WT) vs. endothelial-specific *Lphn2* knockout (ECKO) mice. *En face* examination of descending aortas revealed an approximately 50% reduction in nuclear localization of pSmad1/5 in Lphn2-deficient mice compared to controls (Figure 3A-B). We next examined ECs in zebrafish intersegmental vessels (ISVs). Zebrafish embryos can develop normally for several days without heartbeat or blood flow, allowing assessment of blood flow-dependent signaling without general effects on morphogenesis. Embryos were injected with morpholinos targeting cardiac troponin T (TNNT; *silent heart*) to abolish blood flow ^6,25,26^. Nuclear pSmad1/5 staining in ECs within ISVs of *silent heart* embryos was significantly reduced compared to wild-type embryos. Knockdown of Lphn2 alone produced a similar reduction in pSmad1/5 nuclear signal in ISVs. Notably, combined depletion of TNNT and Lphn2 had no further effect, indicating that Lphn2 and blood flow act within the same pathway (Figure 3C-D). Together, these results confirm a requirement for Lphn2 in flow-dependent activation and nuclear translocation of Smad1/5 in ECs.

**Figure 3:**
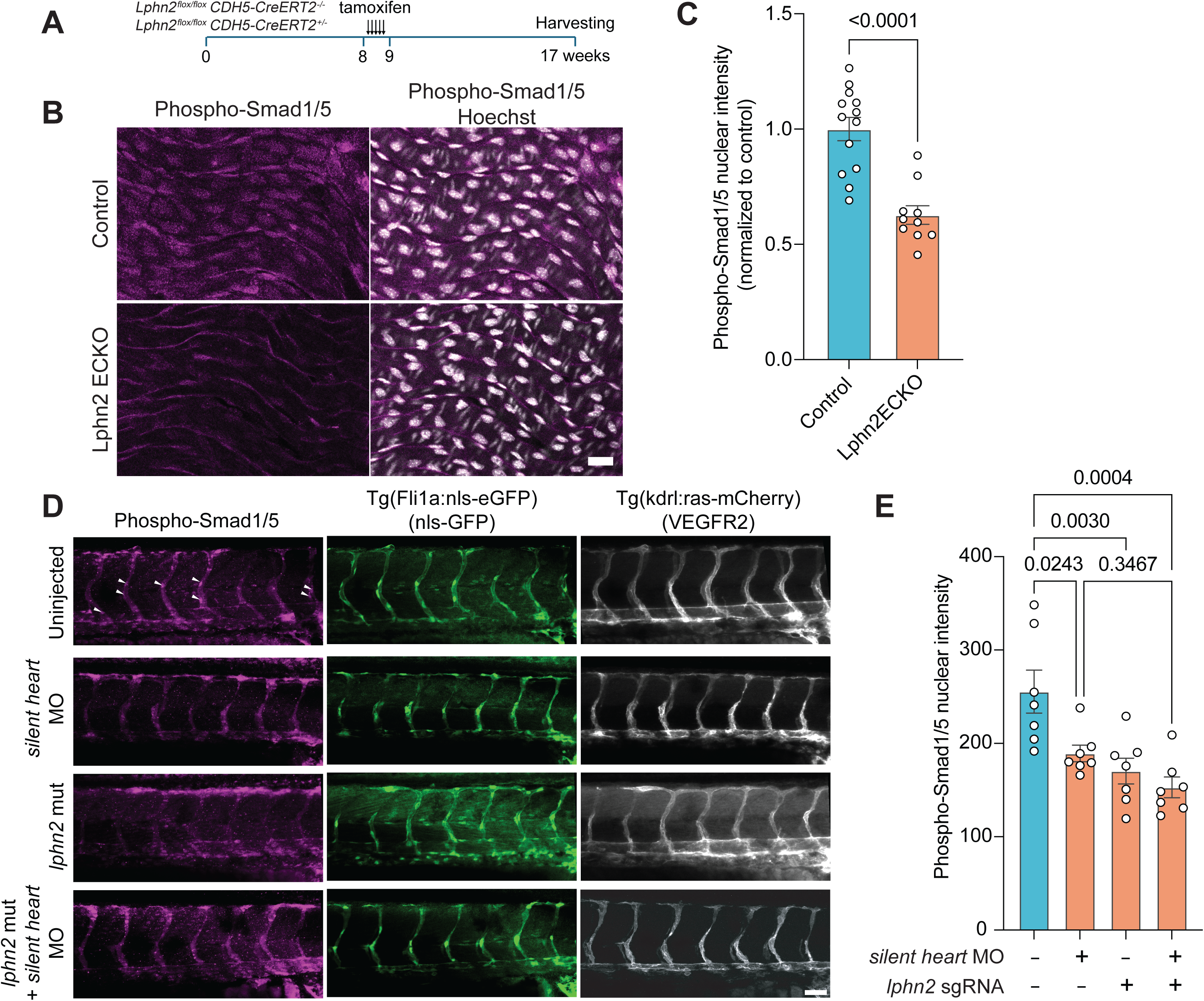
Smad1/5 activity in vivo requires Lphn2. **A**) Experimental timeline: Control or *Lphn2* ECKO mice were injected with tamoxifen at 8 weeks of age and tissue harvested 8 weeks post-injection. **B**) Descending aortas from WT and Lphn2 ECKO mice stained *en face* for phospho-Smad1/5 (pSmad1/5). Scale bar, 50 µm. **C**) Quantification of nuclear pSmad1/5 intensity from (**B**). Values are means ± S.E.M., n = 13 and 10 mice for control and *Lphn2*ECKO, respectively. **D**) Intersegmental vessels in WT and Lphn2 mutant 48 hpf zebrafish embryos stained for pSmad1/5, with or without *silent heart* morpholinos. mcherry-VEGFR2 labels vessels and nls-GFP marks the nuclei. Arrows indicate nuclear pSmad1/5 signal. Scale bar, 50 µm. **E**) Quantification of nuclear pSmad1/5 intensity in (**D**). Values are means ± S.E.M., n = 7 embryos per group. Statistical significance was assessed using Student’s t-test for (**C**) and one-way ANOVA with Tukey post hoc tests for (**E**).

### Plasma membrane fluidization also activates Smad1/5 signaling

Fluid shear stress is reported to reduce plasma membrane cholesterol and increase membrane fluidity (Figure 4A) ^27–30^. We previously used methyl-β-cyclodextrin (MβCD) to reduce membrane cholesterol levels by the same amount as laminar flow, which activated the PECAM1-VEcadherin-VEGFR2 pathway through Lphn2 ^6^. Lphn2 activation thus appears to be regulated by plasma membrane composition or properties. We therefore used MβCD to deplete plasma membrane cholesterol to levels comparable to those induced by 30 minutes of shear stress (Figure 4B). MβCD treatment increased phosphorylation and nuclear localization of Smad1/5 in control siRNA cells, which was abolished by Lphn2 depletion (Figure 4C-F). These findings support a model in which Lphn2 transduces changes in membrane lipid properties into Smad1/5 activation.

**Figure 4:**
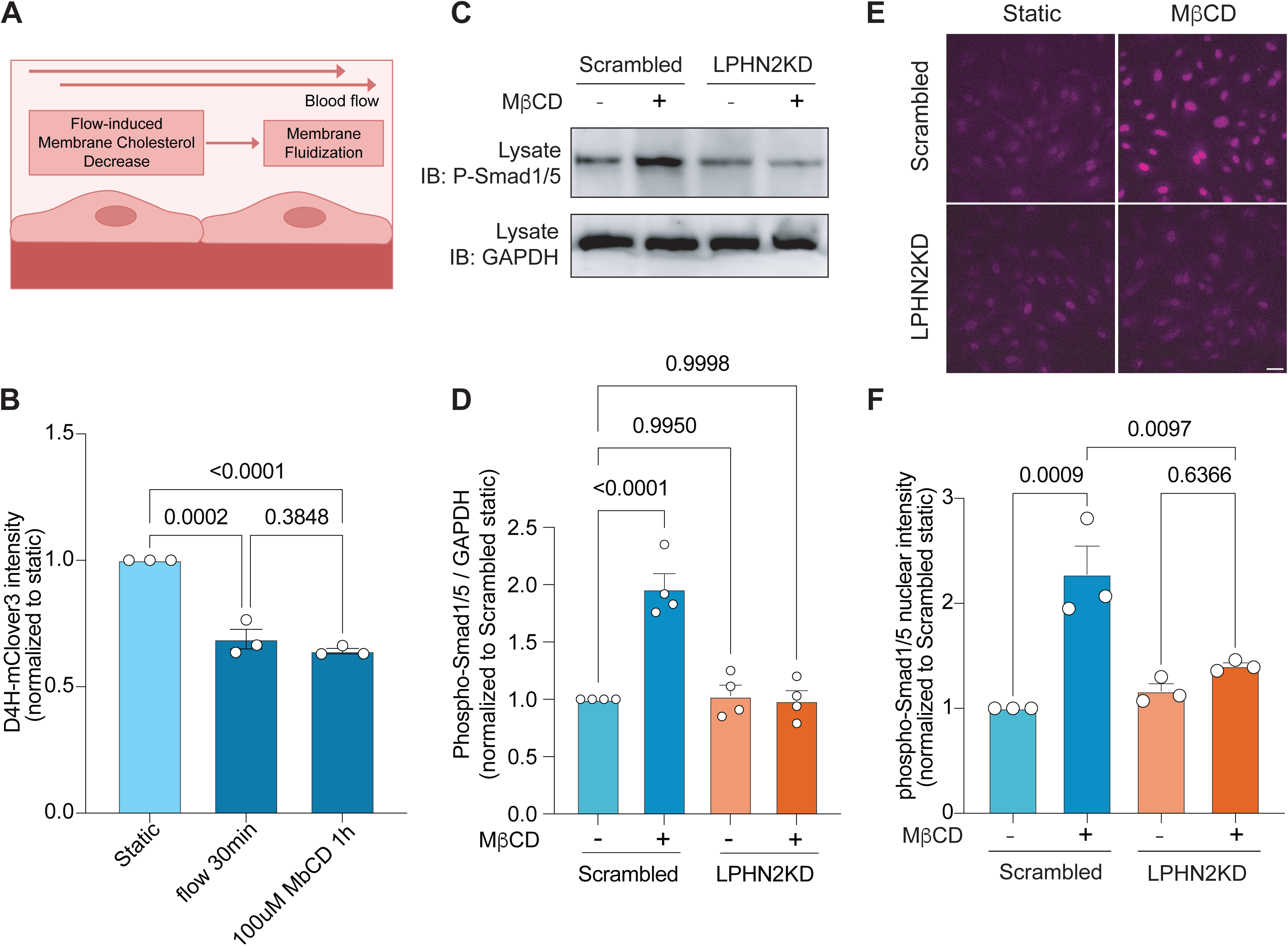
Membrane Fluidization Triggers Smad1/5 Phosphorylation. **A**) Schematic for the connection among blood flow, cholesterol and membrane properties. **B**) Quantification of bound D4H*-mClover3 intensity to confluent HUVEC monolayers. Data are from three independent experiments. Statistics were assessed using one-way ANOVA with Tukey’s multiple-comparisons test. **C**) Phosphorylated Smad1/5 immunoblot in control (scrambled siRNA) and Lphn2 knockdown HUVECs treated with 100 µM methyl-β-cyclodextrin (MβCD) for 1 hour. **D**) Quantification of phosphorylated Smad1/5 normalized to scrambled static (control) in (**C**). Values are means ± S.E.M. from n = 4 independent experiments. **E**) Nuclear localization of phosphorylated Smad1/5 in scrambled siRNA-treated (control) and Lphn2 knockdown HUVECs, treated with 100 µM MβCD for 1 hour, Scale bar, 50 µm. **F**) Quantification of nuclear phosphorylated Smad1/5 intensity in (**E**). Values are normalized to Scrambled static and means ± S.E.M. from three independent experiments. Statistical significance was assessed using one-way ANOVA with Tukey post-hoc tests.

### Lphn2 PDZ domain is required for Smad1/5 activation

We previously found that Eng was dispensable for BMP-induced activation of Smad1/5 but essential for flow enhancement of Smad1/5 activation^8^. Lphn2 harbors a short PDZ-binding motif at its C-terminus, which could conceivably mediate an interaction with Eng via a PDZ domain-containing adapter protein. We therefore investigated whether the Lphn2 PDZ domain was required for its role in the ALK1-Eng-Smad1/5 pathway. Lphn2-depleted ECs were rescued with full-length Lphn2, a GPCR defective mutant, or with two PDZ binding defective Lphn2 mutants-lacking the C-terminal 9 a.a.’s (deltaC9 mutant) or with alanine substitution in the last 3 a.a’s, (3A mutant) (Figure 5A). Nuclear localization of phosphorylated Smad 1/5 nuclear translocation in response to shear stress was then assessed. The GPCR defective mutant was fully functional whereas the PDZ motif-deficient mutants failed to rescue Smad1/5 activation (Figure 5B). The Lphn2 PDZ-binding motif is thus essential for this pathway.

**Figure 5:**
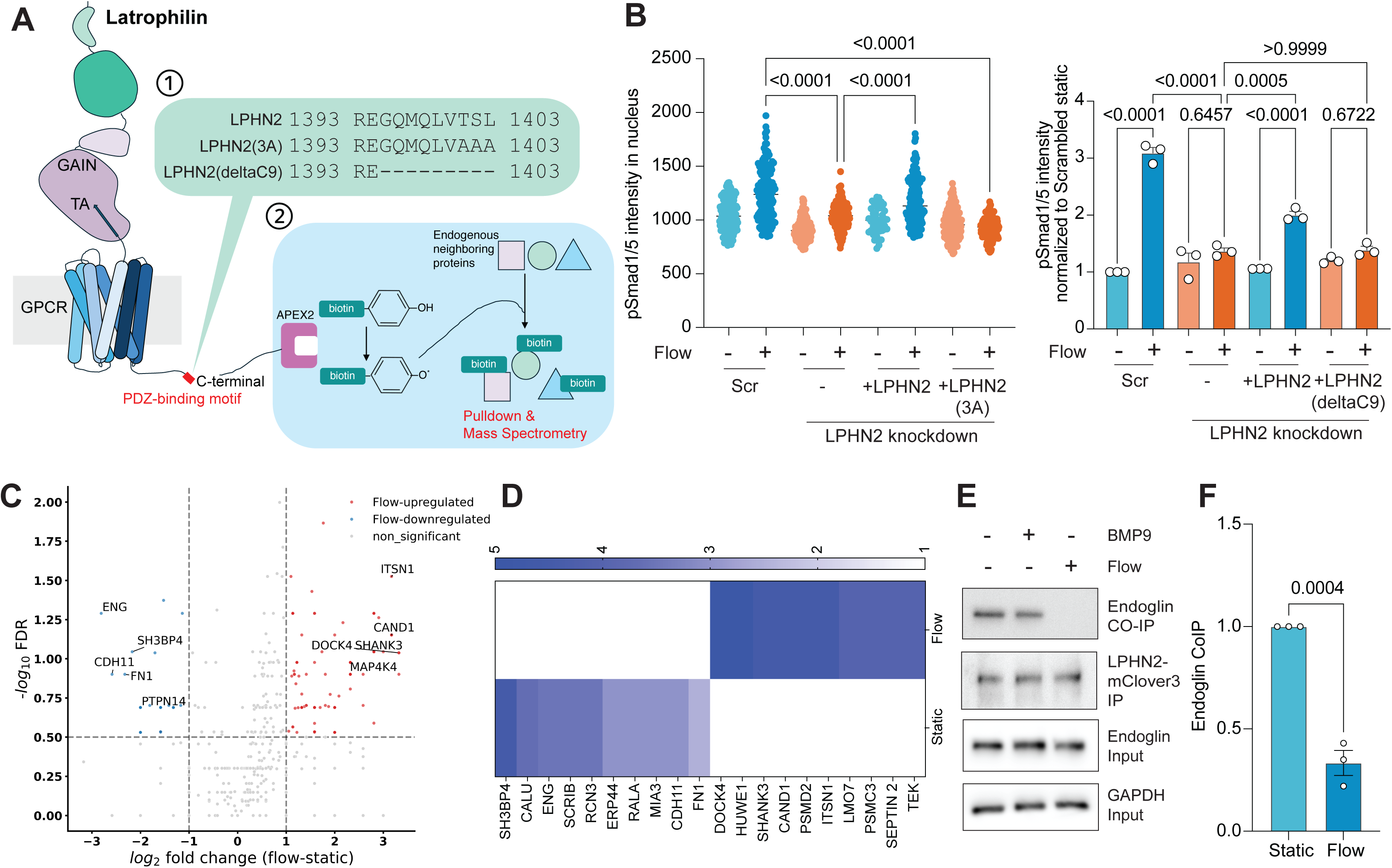
Lphn2 proximity labeling. **A**) Schematic for 1) the sequence of Lphn2 C-terminal PDZ-motif in wild-type (LPHN2) and PDZ-motif mutants with the last three amino acids replaced with alanine-Lphn2 (3A) or the last 9 a.a. deleted (deltaC9) and 2) proximity labeling. **B**) Quantification of nuclear translocation of phosphorylated Smad1/5 in control (Scrambled siRNA/Scr) and Lphn2 knockdown HUVECs rescued with wild-type Lphn2, or the PDZ-binding motif mutants, with or without 30 min of FSS at 12 dynes/cm^2^. Left panel, rescue of Lphn2 KD with PDZ-motif 3A mutant; right panel, rescue with deltaC9 mutant. For left panel: Values are means ± S.E.M. from N = 146, 142, 136, 138, 146, 167, 204, 163 for scrambled static, scrambled flow, Lphn2KD static, Lphn2KD flow, +Lphn2 WT static, +Lphn2 WT flow, +Lphn2(3A) static, +Lphn2(3A) flow, respectively. For right panel-values are means ± S.E.M. from three independent quantifications. Statistical significance was assessed using one-way ANOVA with Tukey post hoc test. **C**) Volcano plot for APEX2-mediated proximity labeling and mass spectrometry-based proteomics, showing differentially enriched proteins associated with Lphn2 under static vs. flow conditions. Five top interactors enriched under only flow or only static are highlighted. **D**) Top 10 interactors of Lphn2 displayed as a heat map based on fold change in peptide counts in flow vs static conditions. **E**) Immunoblot showing co-immunoprecipitation of endogenous endoglin with Lphn2-mClover3 under static conditions, 1 h of laminar flow, or BMP9 (1 ng/mL) stimulation. **F**) Endoglin coimmunoprecipitation normalized to GFP-Lphn2 demonstrated in (**E**). Values are means ± S.E.M. from n = 3 independent experiments. Statistics were assessed using unpaired *Student’s* t-test.

### Interaction with Endoglin

To identify proteins that differentially associate with Lphn2 in a flow-dependent manner, we performed proximity-based proteomics using APEX2-tagged Lphn2, which catalyzes biotinylation of proteins within ∼20 nm in living cells (Figure 5A)^31^. Mass spectrometry of biotinylated proteins identified 182 proteins whose labelling increased in response to flow (Table 1), while 33 proteins decreased (Table 2). Notably, Eng was among the top hits showing reduced labeling after FSS (Figure 5C-D). Co-immunoprecipitation (co-IP) confirmed that Lphn2 and Eng associate at baseline but dissociate in response to flow (Figure 5E and 5F). These findings suggest that Lphn2 interacts with Endoglin, likely requiring its PDZ domain, and regulates its availability for ALK1-mediated Smad1/5 activation, thereby modulating endothelial responses to shear stress.

**Table 1:** List of Lphn2 proximal proteins enriched under flow based on unique peptide counts. 182 proteins with a cut-off of 2 or more unique peptides were identified as flow-specific Lphn2 interactors.

**Table 2:** List of LPHN2 proximal proteins enriched under static conditions based on unique peptide counts. 33 proteins were identified as static-specific interactors of Lphn2 with a cut-off of 2 or more unique peptides.

The interaction between Activin receptor-like kinase 1 (ALK1) and its co-receptor ENG is enhanced by fluid shear stress, which correlates with enhanced Smad1/5 phosphorylation and downstream signaling ^8^. We therefore investigated whether Lphn2 regulates this interaction. Lphn2 depletion strongly reduced the ALK1-Eng association under flow (Figure 6A-B). We also found that Lphn2 associated with Eng and to a lesser extent with ALK1 under static conditions, and this association was disrupted by flow (Figure 6C). The Lphn2-Eng co-IP also required the Lphn2 PDZ domain (Figure 6C). Furthermore, we previously found that flow potentiates BMP9 signaling by shifting the dose response curve toward lower concentrations^8^; we now found that LPHN2 is necessary for this potentiation (Figure 6D-E). Lphn2 thus has a PDZ-dependent interaction with Eng under static conditions, which is disrupted by FSS, possibly releasing Eng to engage ALK1 and facilitate flow-dependent Smad1/5 activation.

**Figure 6:**
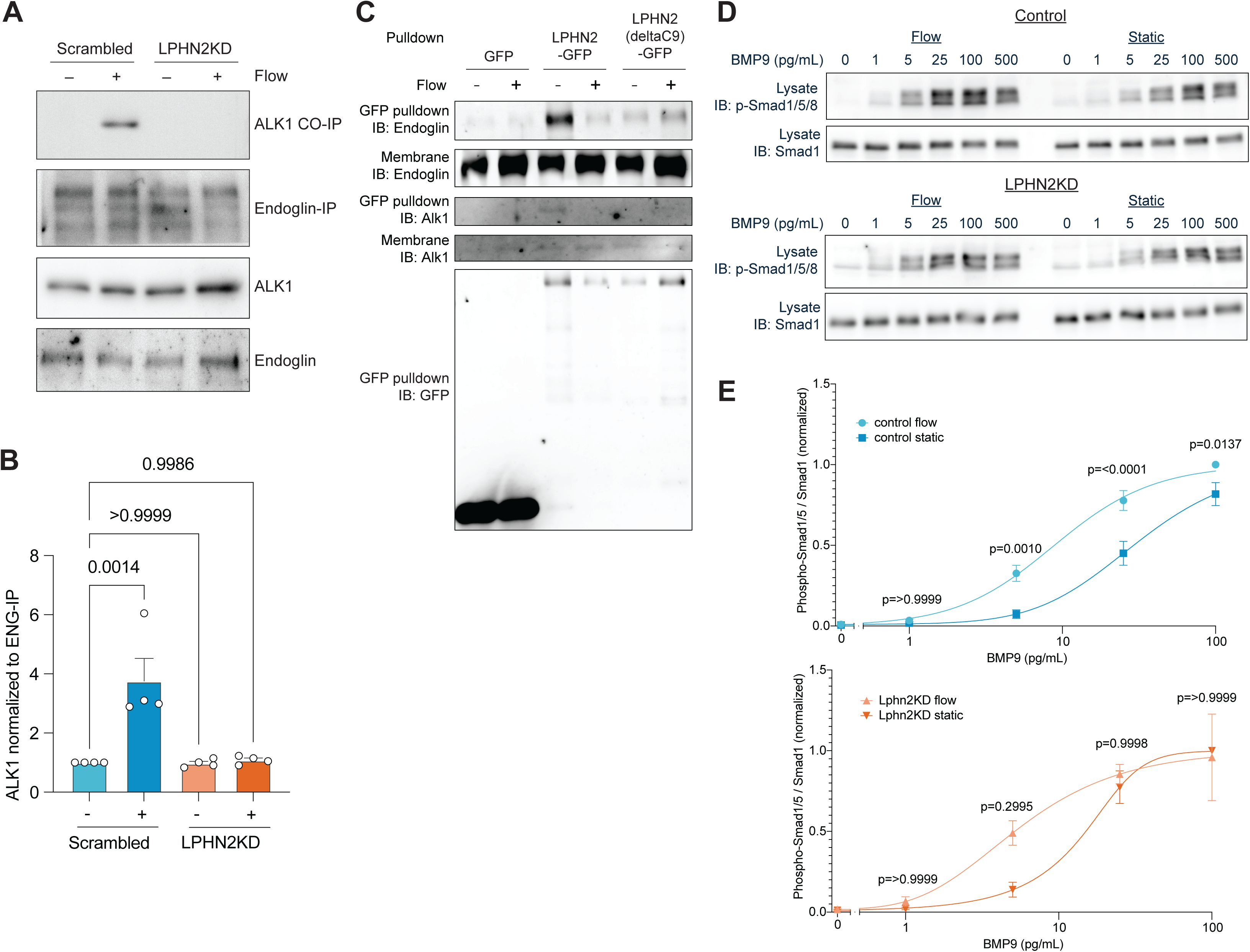
Laminar flow induces Endoglin dissociation from Lphn2. **A**) Co-immunoprecipitation of ALK1 with endoglin in control and Lphn2 knockdown HUVECs with or without 15 minutes of laminar FSS at 12 dynes/cm^2^. **B**) Quantification of ALK1 levels co-immunoprecipitated with endoglin in control and Lphn2-depleted HUVECs with and without FSS. Values are means ± S.E.M. from n=4 independent experiments. Statistical significance was assessed using one-way ANOVA with post-hoc Dunnet test. **C**) immunoblot showing co-immunoprecipitation of endoglin and ALK1 with wild-type Lphn2-GFP or the PDZ-binding-deficient mutant (Lphn2 deltaC9-GFP) with or without 1 hour of laminar flow. **D**) Representative Western blot of Smad1/5 phosphorylation of control or LPHN2KD cells in response to the indicated concentrations of BMP9 in serum-free media with or without flow for 30 minutes. **E**) Quantification of Smad1/5 phosphorylation normalized to total Smad1 from (**D**). Data are from 3-4 independent experiments, and values are mean +/- S.E.M. Statistics used one-way ANOVA with post-hoc Tukey test.

### Interaction with Shank3 in flow-dependent Smad1/5 signaling

Shank3 is a known interactor of the Lphn PDZ binding motif and a well-established component of neuronal synapses, as are Lphns^12^. Shank3 was also one of the top proteins that increased its interaction with Lphn2 upon flow (Fig 5D and Table 1). We therefore investigated its role in Smad1/5 activation by flow and membrane fluidization. Shank3 knockdown blocked activation of Smad1/5 by both FSS and methyl-β-cyclodextrin (Figure 7A-D). Loss of Shank3 also prevented flow-induced release of Eng from Lphn2 (Figure 7E-F) and flow-induced association of Eng and ALK1 (Figure 7G-H). Shank3 depletion also prevented the shift in BMP9 dose-response by flow (Figure 7I-J). To further explore the interaction dynamics among Shank3, Lphn2, and Endoglin in response to flow, we performed co-IP assays using GFP-Shank3, expressed at ∼2X endogenous levels, under static and flow conditions (Supplementary figure 4A-B). Consistent with the proximity proteomics data, flow increased the interaction between Shank3 and Lphn2 while reducing the interaction between Shank3 and Eng (Figure 7K-M). We also tested whether double knockdown of Lphn2 and Shank3 might restore flow-induced Smad1/5 activity, which it did not (Supplemental Figure 5A-B). However, total Endoglin protein was substantially reduced in double knockdown cells (Supplemental Figure 5A and 5C), which likely contributes to decreased flow-induced Smad1/5 signaling. To test the relevance of these results *in vivo*, we examined aortas from global Shank3 knockout mice. Shank3-null mice had reduced EC nuclear pSmad1/5 staining compared to wild-type controls (Figure 8A-B).

**Figure 7:**
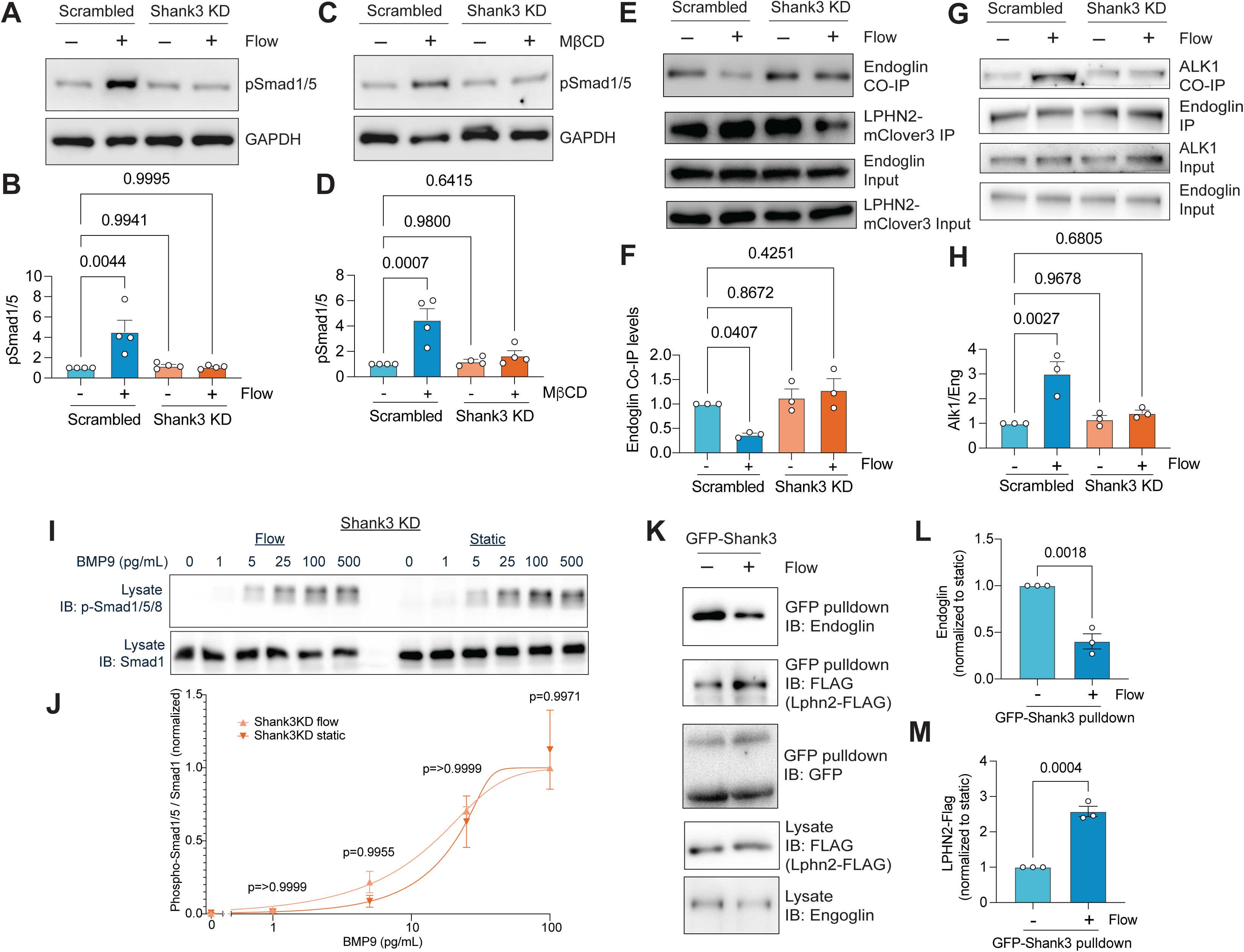
Shank3 in Smad1/5 activation. **A**) Cells were transfected with scrambled siRNA (control) or siRNA targeting *Shank3* with or without 1-hour laminar FSS at 12 dynes/cm^2^. **B**) Quantification of phosphorylated Smad1/5 (pSmad1/5) normalized to scrambled static condition (**A**). Values are means ± S.E.M from n = 4 independent experiments. **C**) Immunoblot of pSmad1/5 in control (scrambled siRNA) and shank3 knockdown HUVECs treated with 100 µM methyl-β-cyclodextrin (MβCD) for 1 hour. **D**) Quantification of phosphorylated Smad1/5 in (**C**) normalized to scrambled static condition. Values are means ± S.E.M. from n = 4 independent experiments **E**) Representative immunoblots of immunoprecipitated Lphn2-mClover3 from HUVECs with or without shank3 knockdown and with or without 1 h laminar flow assessed for co-immunoprecipitated Endoglin. **F**) Quantification of co-IPs from (**E**). Endoglin co-immunoprecipitated with GFP-Lphn2 in control (scrambled siRNA-treated) and *Shank3* knockdown (siShank3) with and without flow. **G**) Co-immunoprecipitation of ALK1 with endoglin in control and *Shank3* knockdown HUVECs with and without 15 minutes of FSS at 12 dynes/cm^2^. **H**) Quantification of ALK1 co-immunoprecipitated with endoglin in control and Lphn2-depleted HUVECs with and without FSS. Values are means ± S.E.M. from n=3 independent experiments. Statistics used one-way ANOVA with post-hoc Tukey test. **I**) Representative Western blot of Smad1/5 phosphorylation of control or Shank3KD cells in response to the indicated concentrations of BMP9 in serum-free media with or without flow for 30 minutes. **J**) Quantification of Smad1/5 phosphorylation normalized to total Smad1 from (**I**). Data are from 3 independent experiments, and values are mean +/- S.E.M. Statistics used one-way ANOVA with post-hoc Tukey test. **K**) Representative immunoblots of immunoprecipitated GFP-tagged Shank3, with and without 15 min laminar flow in HUVECs co-expressing flag-tagged Lphn2 (Lphn2-Flag) and endogenous endoglin. **L&M**) Quantification of Endoglin and Lphn2-Flag co-immunoprecipitated with GFP-Shank3 normalized to static condition. Values are means ± S.E.M. from n = 3. Statistics were assessed using unpaired *Student’s* t-test.

**Figure 8:**
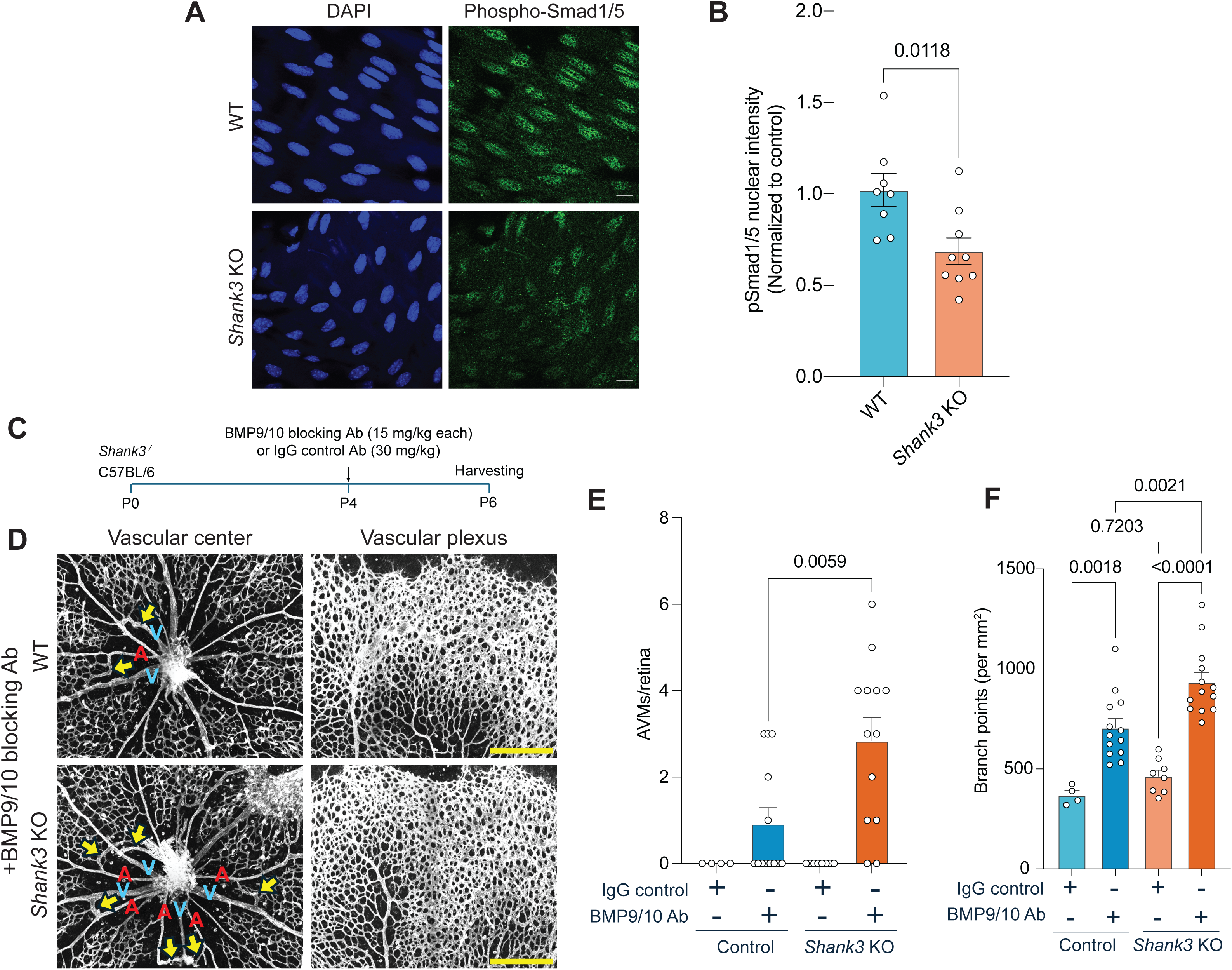
Shank3 KO mice show reduced aortic nuclear pSmad1/5 and increased AVM formation upon BMP9/10 blockade. **A**) Descending aortas from WT and Shank3 global knockout aortas stained *en face* for pSmad1/5 (green) and DAPI (blue) to mark the nuclei. Scale bar, 50 µm. **B**) Quantification of nuclear pSmad1/5 intensity in ECs in (**A**). Values are means ± S.E.M from n = 8 WT and n = 9 Shank3KO mice. **C**) Experimental timeline: Control or *Shank3* KO pups were injected with BMP9/10 blocking antibodies or IgG control antibody at postnatal day 4 (P4), and retinas were harvested at P6. **D**) Representative images of the vascular front (left) and vascular plexus (right) in P6 retinas from pups injected at P4 with BMP9/BMP10-blocking antibodies. V, vein; A, artery. Yellow arrows indicate arteriovenous malformations (AVMs). Scale bar, 500 µm. **E**) Quantification of AVMs at the vascular center in retinas from mice injected with control or BMP9/BMP10-blocking antibodies (n = 13 retinas per group) from (**D**). Values are means ± S.E.M. Statistical significance was calculated using *Student*’s t-test. **F**) Quantification of vascular branch points in Shank3 global KO P6 retinas, normalized by retinal area. Statistics: one-way ANOVA with post-hoc Tukey test.

Impaired ALK1-Endoglin-Smad1/5 signaling leads to arteriovenous malformations (AVMs) in hereditary hemorrhagic telangiectasia^8,10^. We therefore tested whether loss of Shank3 increases susceptibility to AVM formation. To this end, postnatal day 4 (P4) wild-type and Shank3 knockout pups were injected with a moderate dosage of blocking antibodies against BMP9 and BMP10, which is known to mimic HHT (Figure 8C) ^32^. BMP9/10 antibodies induced AVMs near the retinal centers and hypervascularization at the periphery as expected; both of these phenotypes were more severe in Shank3 knockout mice, while no AVMs were seen in IgG-injected retinas from WT or Shank3-null littermates (Figure 8D-8F). Shank3 thus contributes to flow-dependent activation of the ALK1-Endoglin pathway and vascular stability.

As Lphn2 is the upstream receptor for this pathway, we also examined the effect of BMP9/10 antibodies in Lphn2 ECKO mice. These mice showed essentially no AVMs and less hypervascularization after antibody injection (Supplemental Figure 6). However, Lphn2 ECKO retinas with control IgG showed slower vascular expansion. This result is not surprising since AVM progression is known to be flow-dependent^33^, and Lphn2 is required for flow stimulation of angiogenic sprouting^6^. These effects are presumably mediated by other Lphn2-dependent flow pathways, as discussed below.

## Discussion

These data identify the adhesion G protein-coupled receptor Lphn2 as a key mediator of Smad1/5 activation in response to FSS in ECs. Smad1/5 are transcription factors activated downstream of BMP9/10 binding to the ALK1-Endoglin-BMPR2 receptor complex. This signaling pathway promotes vascular quiescence, suppressing excessive sprouting angiogenesis and remodeling ^34,35^. The upstream molecular sensors that link fluid shear stress to the ALK1-Endoglin-Smad axis were, however, previously unknown.

We found that Lphn2 is required specifically for activation of Smad1/5 by FSS but not by BMP9. Surprisingly, this effect was independent of GPCR activity, instead requiring Lphn2’s C-terminal PDZ-binding motif. This sequence likely interacts with scaffolding proteins such as Shank3, which was also required for flow-induced activation of Smad1/5. Co-immunoprecipitation showed that Shank3 associates with both Lphn2 and Endoglin without flow, with flow triggering Shank3 dissociation from Endoglin and association with Lphn2. Concurrently, Endoglin dissociates from junctional Lphn2 and complexes with ALK1 (Supplemental Figure 7). We speculate that Endoglin released from junctional Lphn2 may be positioned to bind a nearby pool of junctional Alk1, forming a high affinity complex that binds circulating or soluble BMP9/10.

FSS reportedly increases plasma membrane fluidity in ECs, associated with a decrease in cholesterol content^27–30^. We previously found that extraction of plasma membrane cholesterol to mimic effects of FSS activated the PECAM pathway through Lphn2 ^6^. Conversely, cholesterol loading of ECs is reported to inhibit responses to flow, which was proposed to contribute to the harmful effects of hypercholesterolemia ^36^. Here, we report that cholesterol depletion also activates Lphn2-dependent Smad1/5 activation. The seven-transmembrane domain spanning Lphn2 seems well positioned to sense changes in membrane properties. Indeed, *in vitro* studies have demonstrated direct effects of membrane composition on GPCR activity^37^. Together, these studies suggest that sensing of membrane properties by the Lphn2 transmembrane domain is important for its activation by FSS in ECs. Elucidating the structural basis by which Lphn2 senses membrane properties and whether tension on its extracellular domain contributes to FSS-induced signaling are additional areas for future studies.

An interesting aspect of the Smad1/5 pathway is its activation by FSS at physiological magnitudes and suppression at high FSS, the latter due to increased expression of BMPER, which inhibits BMP signaling^8,9^. Consistent with vascular stabilization by physiological FSS, several studies have shown that Smad1/5 maintains vascular integrity by inducing genes that suppress cell cycle progression, sprouting angiogenesis and mural cell recruitment^8,9,38^. Conversely, mutations or experimental treatments that block the BMP-Alk1/Eng/Smad1/5 pathway cause hereditary hemorrhagic telangiectasia (HHT) lesions wherein the affected ECs undergo clonal expansion resulting in fragile, leaky malformations with features of high flow-induced outward remodeling^11,39^. Notably, HHT lesions involve activation of PI3K and Akt due to loss of BMP signaling^40^.

The mechanism underlying flow- and Lphn2-dependent regulation of Smad1/5 appears to involve dynamic regulation of protein complexes between the BMP9/10 co-receptor Eng, the type I BMP receptor ALK1, Lphn2 and Shank3. Flow induces release of Eng from Lphn2 and promotes its association with Alk1. Shank3, which contains multiple protein-protein interaction motifs (ankyrin repeats, SH3, PDZ, and SAM domains), also undergoes flow-dependent reorganization of its interactions and is required for the Endoglin switch from Lphn2 to Alk1. These data thus suggest a complex mode of regulation. Further analysis of these protein complexes and their regulation by flow will be required to fully understand the control of Smad1/5 activation in ECs.

The increased susceptibility of Shank3 knockout mice to AVM formation after BMP9/10 inhibition highlights a role for junctional mechanotransduction in safeguarding vascular patterning. While mutations in ALK1, Endoglin or (more rarely) Smad4 are required for AVM formation in HHT, only a small subset of affected vessels form AVMs, indicating that additional stresses or hits are required for pathological remodeling. Shank3 deletion alone did not induce spontaneous AVMs but significantly worsened vascular malformations when BMP9/10 signaling was inhibited, supporting the idea that components of the Lphn2-Shank3 pathway may be among the additional hits that contribute to disease. Such mutations or conditions might explain the wide range of symptoms in patients with similar mutations^41,42^.

In an ostensibly contradictory result, Lphn2 ECKO did not induce AVMs, indeed, when combined with injection of BMP9/10 antibodies, AVMs were reduced. But as noted above, HHT lesions require additional activating events, particularly increased PI3K/Akt activity^43,44^. Flow-induced PI3K activation requires PECAM^11,45^, which is downstream of Lphn2^6^. Thus, simultaneous loss of both Smad1/5 and PI3K-Akt with Lphn2 ECKO can explain the absence of AVMs after deletion of Lphn2.

The accompanying submission identifies Lphn2 as the upstream mediator of a third flow-dependent pathway, Notch activation. These results position Lphn2 as a central mechanosensory hub in the endothelium, coordinating biophysical inputs and scaffolding dynamics to regulate three downstream pathways. Importantly, different Lphn2 mutants in these papers selectively block each of the three downstream pathways, arguing that they are separable. Available data suggest that signal integration occurs mainly at the level of gene expression. For example, Notch and Smad1/5 coordinately regulate expression of genes that control vascular stability^46^. Conversely, Smad1/5 regulate genes that restrain PI3K activation by flow^9,43^. A deeper understanding of how these downstream pathways coordinately regulate vascular remodeling and stability under different conditions is a promising direction for future research that is likely to reveal novel therapeutic targets for treating a wide range of vascular diseases.

## Methods

### Plasmids, Cell Culture and siRNA Transfection

Lphn2 with C-terminal mClover3 was described in ^6^. Lphn2-H1071A mClover3 and Lphn2 PDZ-binding motif mutants with last 3 amino acids substituted with alanine (3A mutant) and mutant lacking the last 9 amino acids (deltaC9) were generated using a site-directed mutagenesis kit (New England Biolabs) as per the manufacturer’s instructions. Lphn2-APEX2 was generated by replacing mClover3 with FLAG-tagged APEX2. APEX2 was first cloned into the pBob-GFP vector by substituting GFP with FLAG-tagged APEX2 using Gibson assembly. This process included introducing a silent mutation in APEX2 to eliminate the AgeI site, enabling the utilization of the AgeI site in the pBob lentiviral vector backbone. Subsequently, Lphn2 was transferred from the Lphn2-mClover3 through restriction enzyme digestion and DNA ligation. GFP-tagged Shank3 was a kind gift from Dr. Yong-Hui Jiang (Yale University). All the Lphn2 constructs used in this study were designed to be resistant to Lphn2 siRNAs (Horizon Discovery; L-005651-00-0005)

Primary Human umbilical vein endothelial cells (HUVECs) were obtained from the Yale Vascular Biology and Therapeutics core facility. Each batch contains cells pooled from three donors. Cells were cultured in M199 (Gibco: 11150-059) supplemented with 20% FBS, 1× penicillin-streptomycin (Gibco:15140-122), 60 μg/ml heparin (Sigma: H3393), and endothelial growth cell supplement (hereafter, complete medium). HUVECs were used for experiments at passages 3-6.

For knockdowns, siRNAs against human Lphn1 (Horizon Discovery; L-005650-00-0005), Lphn2 (Horizon Discovery; L-005651-00-0005), Lphn3 (Horizon Discovery; L-005652-00-0005), Gai, Gq/11 ^47^ ARRB1 and ARRB2 (Horizon Discovery-L-011971-00-0005 and L-007292-00-0005) and human SHANK3 (Horizon Discovery; L-024645-00-0005) were used as described^6^. Briefly, 20 nM of siRNAs against Lphn2 and Shank3 were mixed with Lipofectamine RNAi max and added to HUVECs for 4 h in Opti-MEM. Cells were then incubated in EGM-2 (Lonza; CC-3162) for 48 h. For experiments, cells were re-seeded in M199 media.

### Fluid shear stress experiments and MβCD treatment

Laminar shear stress was applied using parallel flow chambers as described ^6,48^. HUVECs on glass slides or tissue culture-treated plastic slides coated with fibronectin (10 μg/ml) for 1 h at 37 °C and grown to confluence in HUVEC complete media were serum-starved in 2% FBS for overnight and placed in parallel flow chambers and shear stress at 12 dynes/cm^2^ applied for the indicated times, quickly washed in 1x PBS, pH 7.4 and lysed in 1x Laemmli buffer.

For cholesterol depletion, HUVECs were incubated in M199 medium containing 100 µM methyl-beta-cyclodextrin (MβCD) and 20 pg/mL of BMP9 for 1 hour. Control cells were treated with M199 with 20 pg/mL of BMP9 for 1 hour. Cells were quickly washed in PBS, pH 7.4 and scraped in 1x Laemmli buffer.

### Immunoblotting and Immunofluorescence for Smad1/5 Activation

Protein lysates were resolved by SDS-PAGE and transferred to nitrocellulose membranes. Membranes were blocked with 5% skim milk in Tris-buffered saline, pH 7.4 containing 0.1% Tween-20 and incubated with antibodies against phosphorylated Smad1/5 (Cell Signaling Technology; 13820), total Smad1 (Cell Signaling Technology, 9743), Gi (NewEast Biosciences, 26003), Gq11 (BD Bioscience, 612705), tubulin (Invitrogen, 62204) and GAPDH (Cell Signaling Technology; 2118). For imaging experiments, cells were fixed in 4% paraformaldehyde following exposure to flow either in the flow chamber or on an orbital shaker, permeabilized with phosphate buffered saline (PBS), pH 7.4 containing 0.1% Triton-X100 for 10 mins and incubated with antibody against pSmad1/5 (Cell Signaling Technology; 9516) overnight at 4°C, followed by incubation with fluorescently labelled secondary antibodies and Hoechst to label nuclei. Images were acquired on PerkinElmer spinning disk using a 20x objective. Nuclear pSmad1/5 was quantified by a custom ImageJ macro code that creates nucleus masks derived from Hoechst-stained images and quantifies the intensity of phospho-Smad1/5 images within the nuclei.

### RNA-sequencing sample preparation and data analysis

RNA was extracted from HUVECs with control or Lphn2 siRNAs using the QIAGEN RNeasy kit according to the manufacturer’s instructions. RNA-sequencing libraries were prepared using poly-A enrichment and sequenced at the Yale Center for Genomic Analysis.

Raw sequencing reads were checked for quality with FastQC (v0.12.1). Adapter sequences and low-quality bases were removed using fastp (v0.23.2). Clean reads were aligned to the human reference genome (GRCh38) using STAR (v2.7.11a). PCR duplicates were removed with Picard MarkDuplicates (v2.25.6). Gene-level read counts were generated using featureCounts from the Subread package (v2.0.3) with GENCODE annotations. Batch effects were corrected using ComBat-seq from the sva package (v3.56.0), and counts were normalized using edgeR (v4.6.2).

### Mouse and Zebrafish Models experiments

The vascular system of Lphn2 ECKO mice^6^, Shank3 global knockout mice^49^ and their littermate controls were perfused with PBS followed by 4% paraformaldehyde (PFA). Aortas were harvested, opened longitudinally, and processed for *en face* immunostaining. Tissues were stained with anti-phospho-Smad1/5 antibody (CST; 9516) and DAPI, and nuclear pSmad1/5 signal was quantified in the descending aorta by manually making masks on EC nuclei using Image J. For transverse section of the aortas, staining was performed as described above using pSmad1/5 antibody (Millipore sigma AB3848-I) and anti-CD31 antibody (BD Science; 557355) to stain the endothelium. pSmad1/5 intensity normalized to DAPI intensity within CD31 positive endothelium was quantified using image J.

To examine the contribution of blood flow to Lphn2-dependent Smad1/5 activation, we utilized zebrafish (*Danio rerio*) embryos, which develop normal vasculature in the absence of a heartbeat for the first few days’ post-fertilization. To inhibit blood flow and assess the requirement of Lphn2, zebrafish embryos at the one-cell stage were injected with a control morpholino or 1 nL of 1 µg/µL of translation-blocking morpholino targeting cardiac troponin T (tnnt2a) (silent heart phenotype) together with Lphn2 sgRNAs or control sgRNAs, and an equivalent concentration of Cas9 endonuclease ^6,25,26^. At 48 hours post-fertilization, embryos were fixed in 4% PFA and stained for pSmad1/5 (Cell signaling, cat #:13820) followed by AlexaFluor 546-conjugated anti-Rabbit 2^nd^ antibody. Nuclear localization of pSmad1/5 in endothelial cells of the intersegmental vessels (ISVs) was assessed by confocal microscopy and quantified using ImageJ.

AVM formation in mouse retina was performed as described^32^. For Lphn2 ECKO pups, tamoxifen (100mg/kg) were administered intragastrically on postnatal days 1, 2 and 3. On postnatal day 4 (P4), mice received BMP9- and BMP10-blocking antibodies (MAB3209 and MAB2926, R&D systems) or IgG-control antibody (MAB003, R&D systems) by intraperitoneal injection of 15 mg/kg each for blocking antibodies or 30 mg/kg for IgG control antibody. Eyes from pups were collected at P6 for the Shank3 KO experiment or at P7 for the Lphn2 ECKO experiment, fixed in 4% paraformaldehyde for 20 min at room temperature, and then dissected to isolate retinas. Isolated retinas were permeabilized and blocked for 1 h at room temperature in blocking buffer containing 1% fetal bovine serum, 3% bovine serum albumin, 0.5% Triton X-100, and 0.01% sodium deoxycholate in PBS (pH 7.4). Retinas were then incubated with isolectin B4 (I32450; ThermoScientific) and indicated fluorescent secondary antibodies for 1 h at room temperature. After washing, samples were mounted in fluorescent mounting medium (DAKO, USA). Confocal images were acquired using a Leica SP8 confocal microscope equipped with a spectral detection system (15 SP detector; Leica Biosystems). Quantitation of retinal vascular parameters was performed using AngioTool or built-in skeletonization function of ImageJ/Fiji, and arteriovenous malformations were quantified in a blinded manner.

### Proximity Labeling and Mass Spectrometry Analysis of Lphn2 Interactome

HUVECs were transduced with lentivirus encoding Lphn2-APEX2 fusion protein or membrane-localized APEX2 and selected for uniform expression. Cells were left in static medium or exposed to laminar shear stress (12 dynes/cm²) for 24 hours. To induce biotinylation, cells were incubated with 2.5 mM biotin-phenol (Iris Biotech) for 30 minutes, followed by rapid addition of 1 mM hydrogen peroxide (H₂O₂) for 1 minute to initiate labeling. The reaction was quenched with Trolox, sodium ascorbate, and catalase, and cells were immediately lysed in RIPA buffer supplemented with protease inhibitors. Biotinylated proteins were enriched using streptavidin-conjugated magnetic beads (Thermo Scientific), washed twice with RIPA buffer, then once with 1M KCl, once with 0.1M sodium carboxylate, once with 2M urea in Tris-HCl (pH7.5) and twice with RIPA again. Proteins were eluted by boiling in 2x Laemmli buffer containing 2mM biotin and 20mM DTT. Proteins were loaded on SDS-PAGE, run for a short time until they just entered the running gel, and the gel was stained with Coomassie blue. The gel pieces containing the proteins were excised and sent to the Taplin Biological Mass Spectrometry Facility (TMSF), Harvard Medical School, Boston, USA. Following LC-MS/MS experiment, proteomics data were analyzed using MaxQuant and visualized using the Perseus platform^50,51^.

### Immunoprecipitation

HUVECs were transduced with lentiviral particles containing epitope-tagged constructs (Lphn2-mClover3 full-length or Lphn2-PDZ domain mutant or GFP-Shank3) for 48 hours and maintained in static conditions or exposed to laminar shear stress (12 dynes/cm²) for indicated times. Cells were lysed in ice-cold NP-40 lysis buffer (50 mM Tris-HCl pH 7.5, 150 mM NaCl, 1% NP-40, 1 mM EDTA) supplemented with protease and phosphatase inhibitors. Lysates were clarified by centrifugation at 14,000 × g for 15 minutes at 4°C. Equal amounts of protein lysates were precipitated using GFP-Trap beads (Chromtek; gta) for 4 hours at 4°C with end-over-end mixing. Beads were washed four times with lysis buffer, and bound proteins eluted by boiling in SDS sample buffer. Samples were resolved by SDS-PAGE and analyzed by immunoblotting using antibodies against Endoglin (Biotechne R& D systems; AF1097), Alk1 (Biotechne R& D systems; AF370), GFP (Thermofisher scientific; A-11122) and Flag (Cell Signaling Technology; 14793). Input controls (5-10% of total lysate) were included in all experiments. For the confirmation of GFP-Shank3 expression, Shank3 antibody (Cell Signaling Technology; 64555) and Vinculin antibody (Sigma; V9131) were used.

### Statistical analysis

All quantitative data are presented as means ± standard error of the mean (S.E.M.), unless otherwise specified. Comparisons between two groups were performed using a two-tailed unpaired Student’s *t*-test. For comparisons involving more than two groups or conditions, one-way analysis of variance (ANOVA) followed by appropriate post hoc tests was used to determine statistical significance. For cell-based quantifications, individual cells were counted across at least three independent experiments, with sample sizes (*N*) indicated in the corresponding figure legends. Protein expression levels from immunoblots were quantified by densitometry and normalized to loading controls such as total Smad1 or tubulin. For proximity labeling, statistical significance for differential protein enrichment between conditions (static vs. flow) was assessed using mass spectrometry-derived abundance values and plotted as volcano plots, with thresholds for significance typically set at *p* < 0.05 and fold change ≥1.5, unless otherwise stated. All statistical analyses were performed using GraphPad Prism (version 9.5.1) and *p*-values were calculated as indicated in the figure legends.

## Supplementary figures

**Supplementary figure 1:**
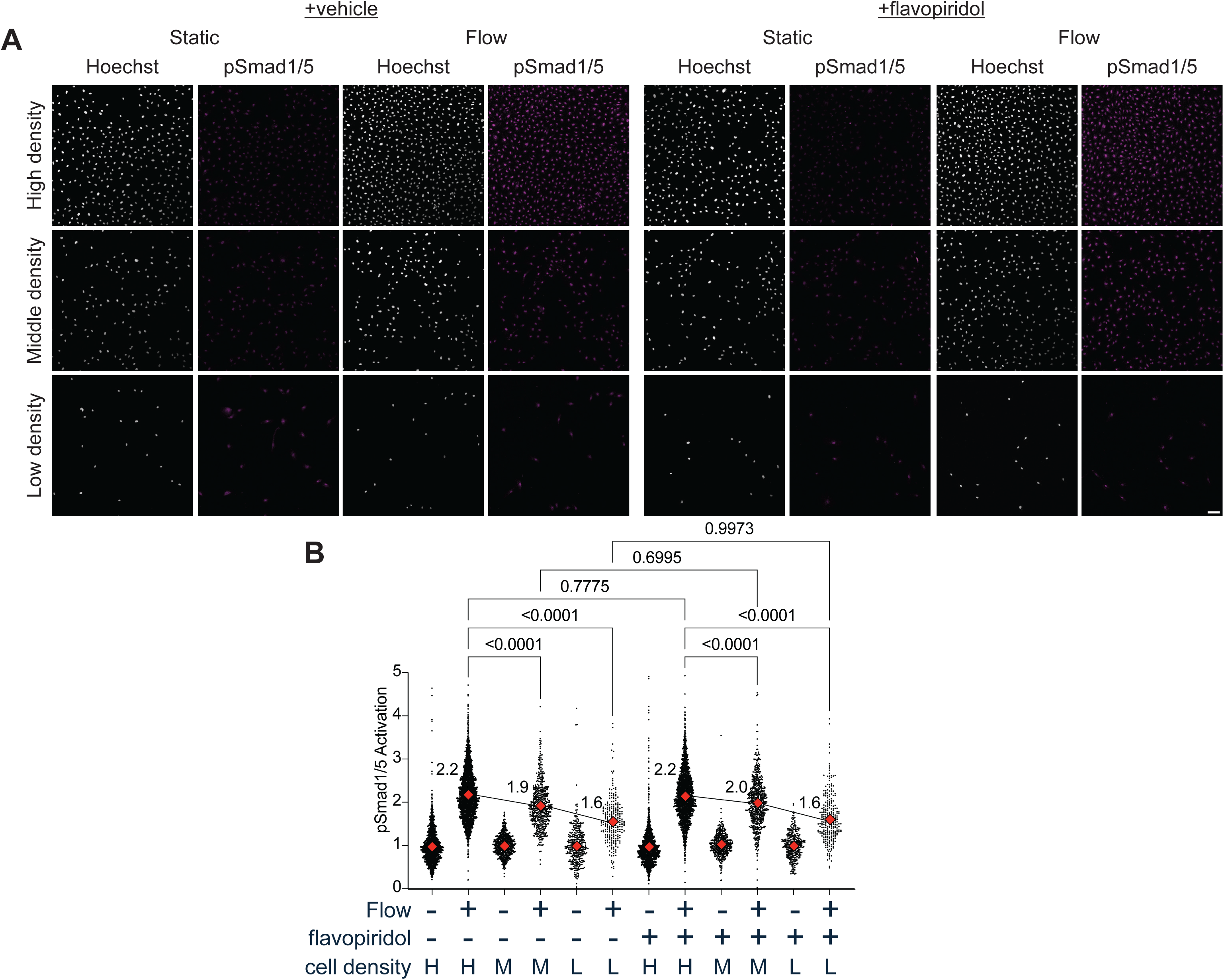
Cell-cycle blockade does not affect density-dependent Smad1/5 activation. **A**) Representative images showing nuclear localization of phospho-Smad1/5 in HUVECs seeded at different densities, with or without 30 min of flow, in the presence or absence of pan-CDK inhibitor flavopiridol (0.2µM, for 1 hour total, added 30 minutes prior to flow onset). **B**) Quantification of pSmad1/5 nuclear intensity from (**A**). N= 1403, 2186, 656, 514, 344, 255, 1365, 1961, 410, 590, 329, 262 cells from left to right as shown in the graph, respectively. For cell density, H, M, L indicates high, medium and low density, respectively. Statistics were assessed using one-way ANOVA with Tukey post hoc tests.

**Supplementary figure 2:**
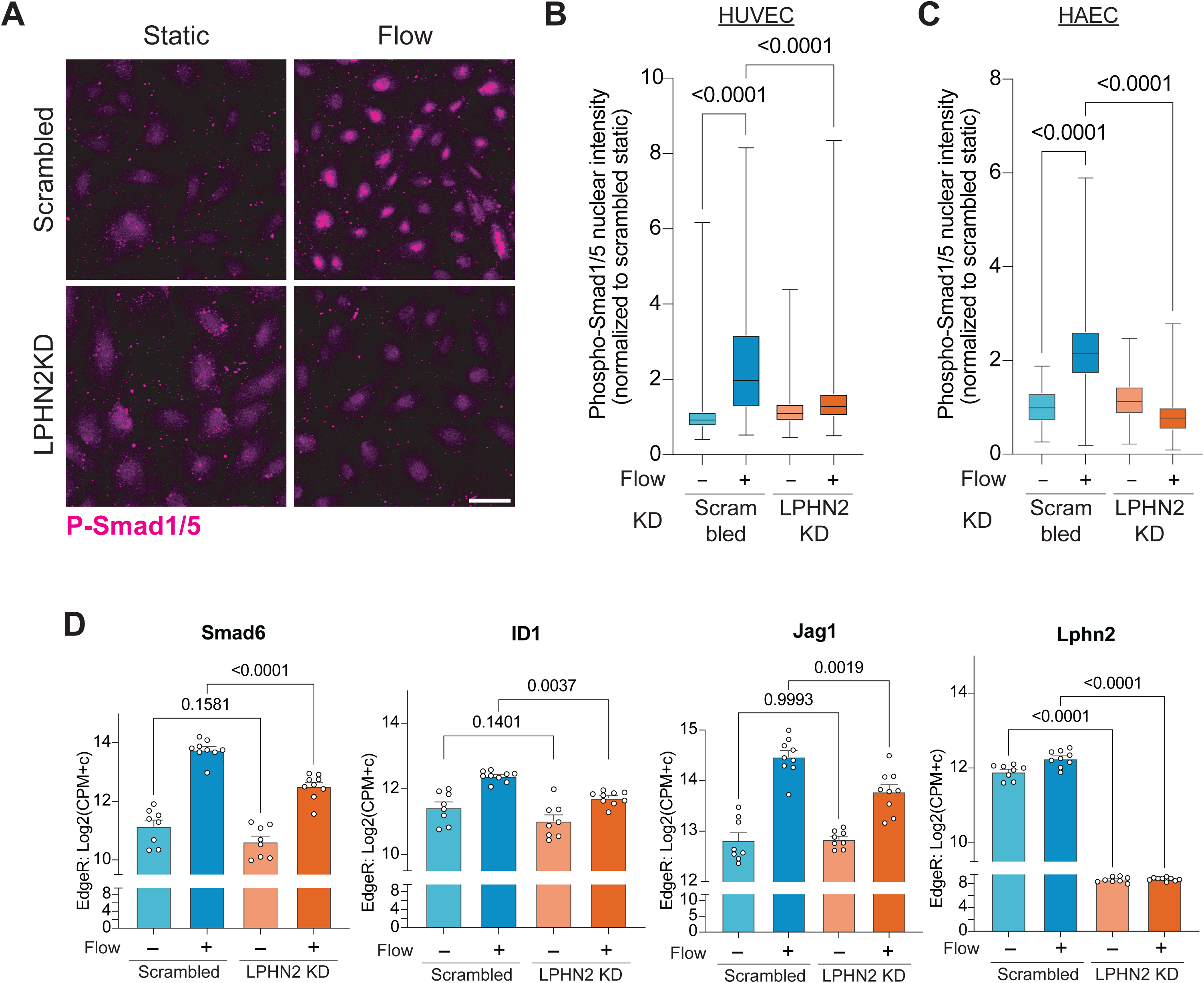
Further validations of Lphn2-dependent flow-induced Smad1/5 signaling. **A**) Representative images showing nuclear localization of pSmad1/5 in HUVECs treated with scrambled and LPHN2 siRNA with and without FSS. Quantification of pSmad 1/5 nuclear intensity upon LPHN2 depletion in HUVECs (**B**) and HAECs (**C**) with and without FSS. Data are presented as box-and-whisker plots showing median (center line), interquartile range (box), and min–max whiskers. N= 717, 1356, 658, 1085 HUVECs and N= 51, 308, 314, 476 HAECs for scrambled static, scrambled flow, LPHN2 KD static and LPHN2KD flow, respectively. Statistics were assessed using one-way ANOVA with Tukey post-hoc tests. (**D**) Normalized expression for the canonical Smad1/5 targets, including *Smad6*, *Id1* and *Jag1,* and *Lphn2* in bulk RNA-sequencing samples treated with control or *Lphn2* siRNA, with or without flow, analyzed by edgeR. Fold changes (flow vs. static) were as follows: *Smad6* (6.1-fold scrambled; 3.7-fold LPHN2 KD), *Id1* (1.9-fold scrambled; 1.6-fold LPHN2 KD), *Jag1* (3.2-fold scrambled; 1.9-fold LPHN2 KD), *Lphn2* (1.3-fold scrambled; 1.1-fold LPHN2 KD). Statistical significance was assessed by one-way ANOVA with Tukey post-hoc analysis.

**Supplementary figure 3:**
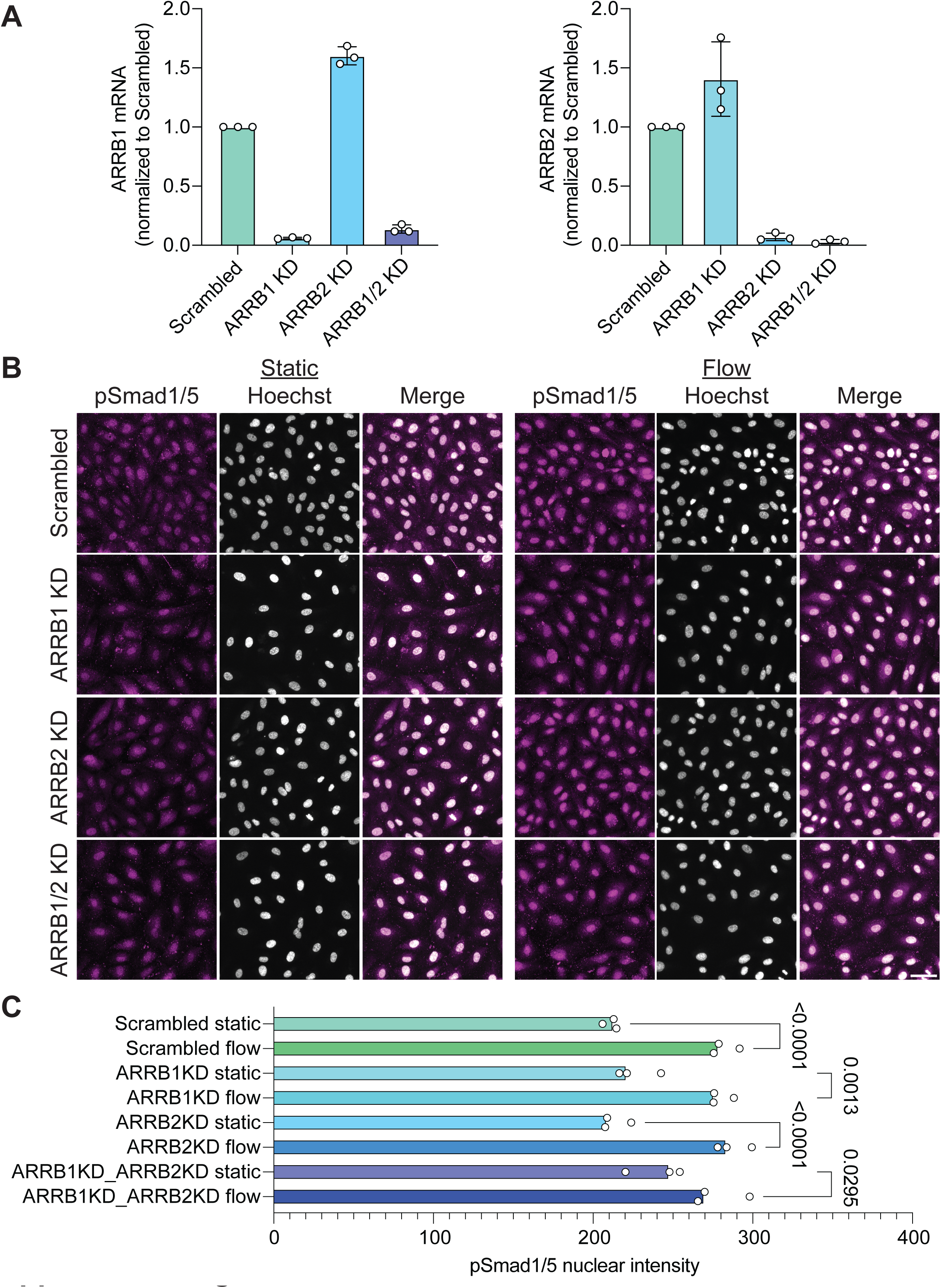
β-arrestins are not required for flow-induced Smad1/5 activation. **A**) mRNA levels of β-arrestins, ARRB1 (left panel) and ARRB2 (right panel) in HUVECs treated with scrambled siRNA, ARRB1 siRNA, ARRB2 siRNA and combined ARRB1 and ARRB2 siRNA (ARRB1/2) measured by quantitative PCR and expressed as fold change relative to scrambled control. Values are means ± S.E.M. from n = 3 independent experiments. **B**) Representative images showing pSmad1/5 (magenta) and Hoechst-stained nuclei (gray) in HUVECs subject to control, ARRB1, ARRB2 and combined ARRB1 and ARRB2 knockdown (ARRB1/2), with and without flow (static). Scale bar, 50 µm. **C**) Quantification of nuclear pSmad1/5 levels in (**B**). Values are means ± S.E.M. from three independent experiments. Statistics were assessed using one-way ANOVA with Tukey post hoc tests.

**Supplementary figure 4:**
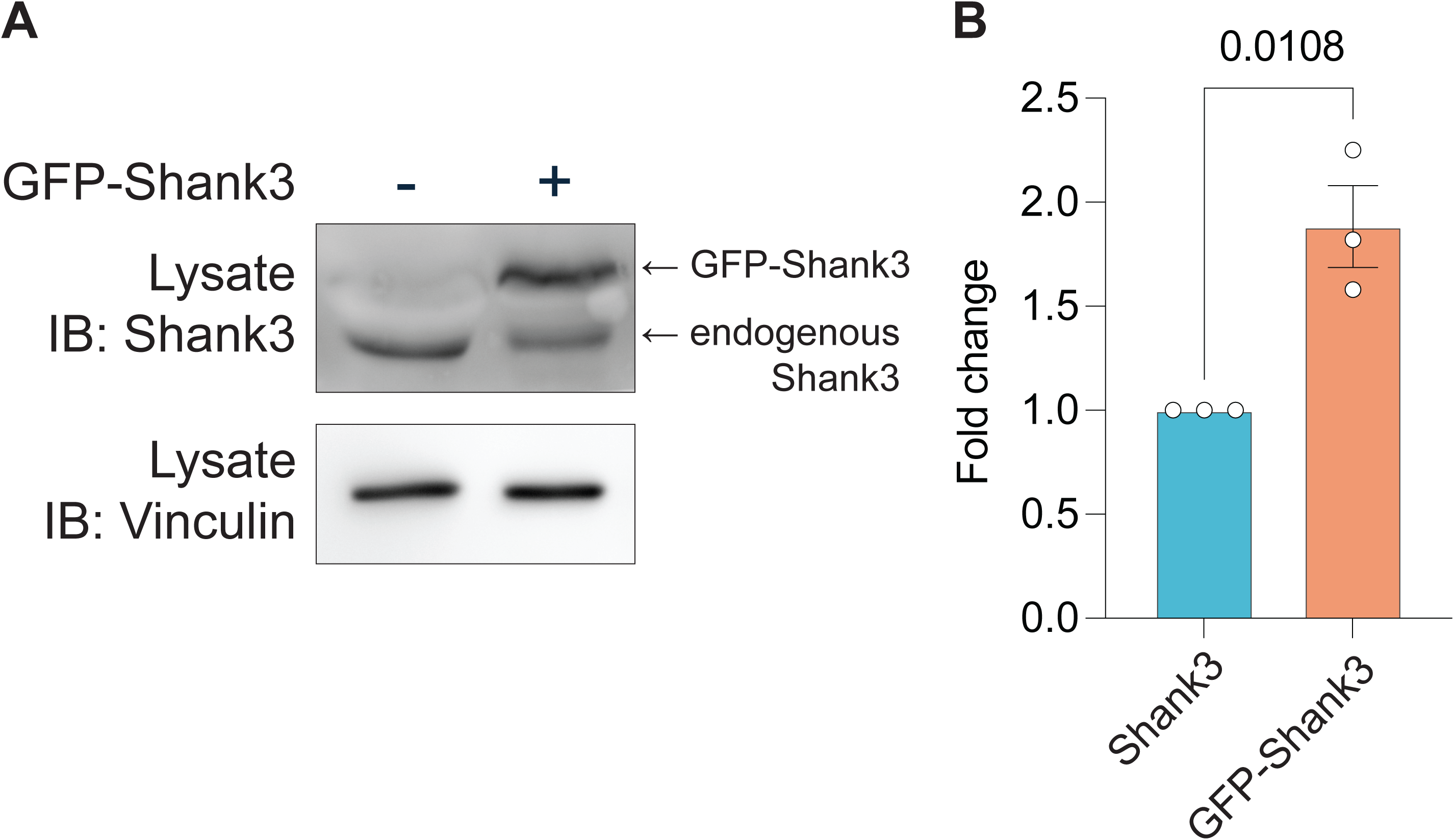
Expression validation of GFP-Shank3. **A**) Representative immunoblot and quantification (**B**) of Shank3 levels in control and GFP-Shank3 over-expressing HUVECs.

**Supplementary figure 5:**
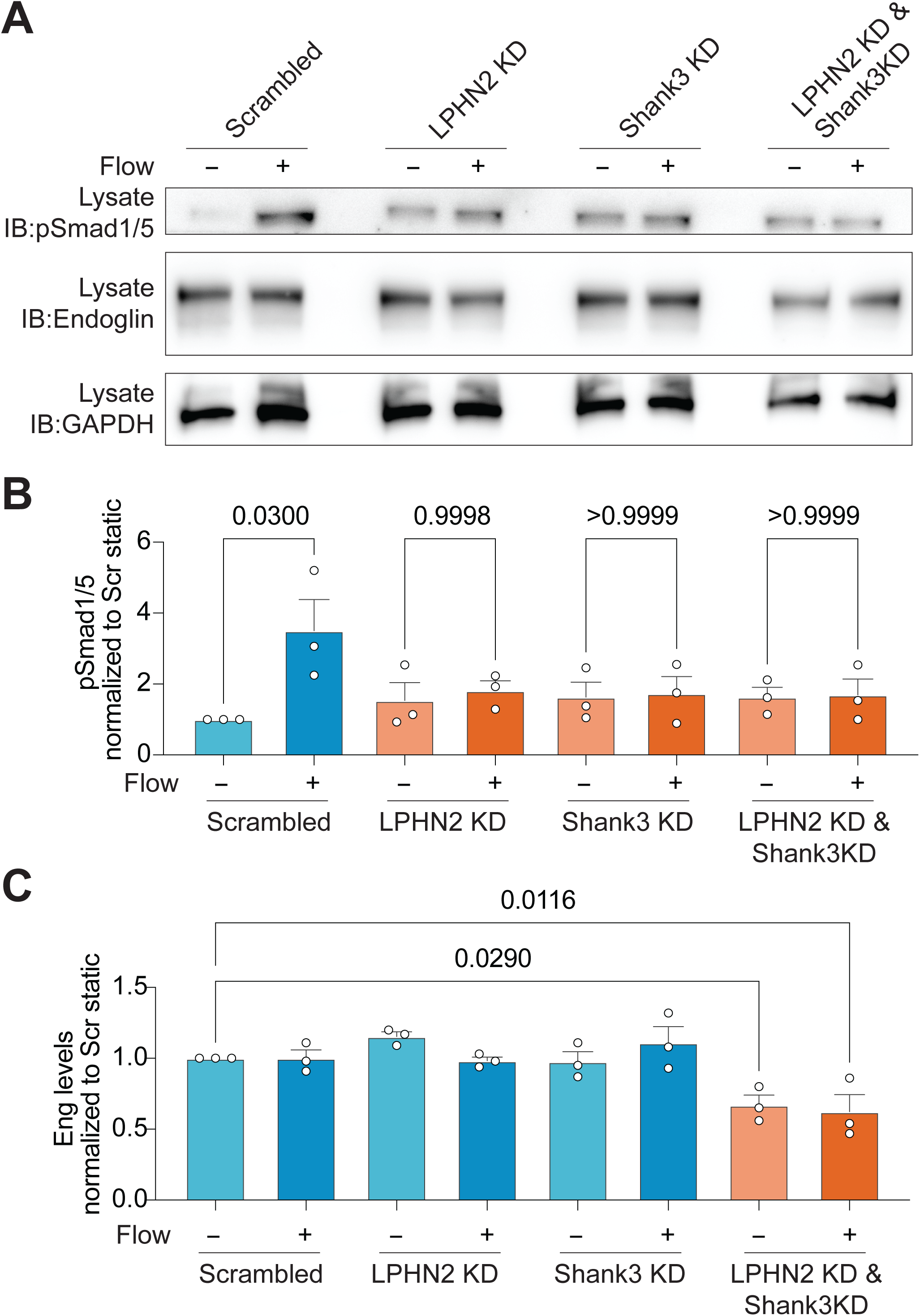
Effect of LPHN2 and SHANK3 knockdown on Smad1/5 activation and Endoglin levels. **A**) Representative Western blot for Smad1/5 phosphorylation and Endoglin expression levels, with or without flow, under control, LPHN2 KD, Shank3 KD and double KD conditions. **B**) Quantification of Smad1/5 phosphorylation from (**A**). Data are from three independent experiments. Statistics used one-way ANOVA with post-hoc Tukey test. **C**) Quantification of Endoglin level from (**A**). Statistic used one-way ANOVA with post-hoc Tukey test.

**Supplementary figure 6:**
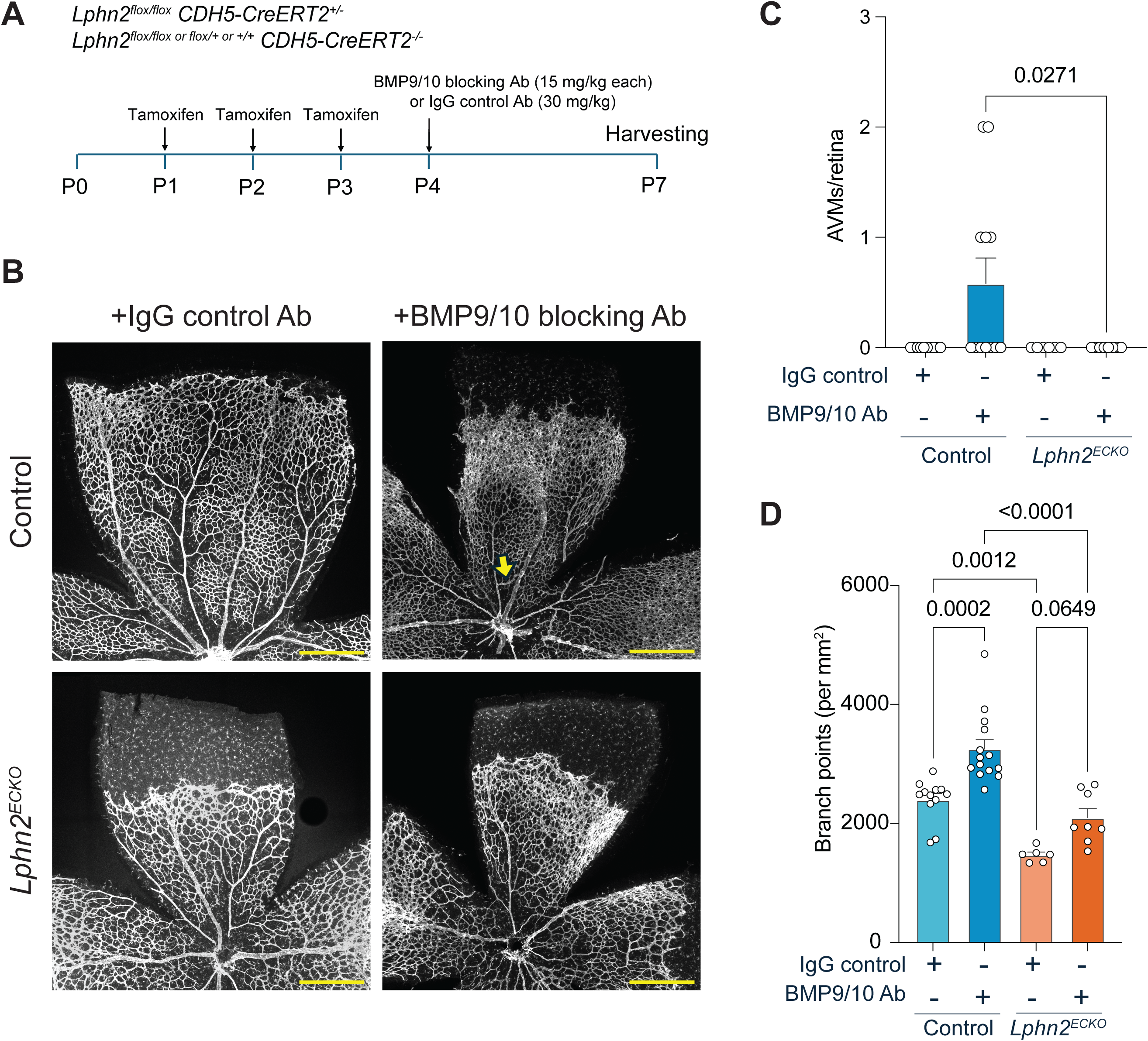
Lphn2 ECKO blocks AVMs and vascular overgrowth induced by BMP9/10 blockade. **A**) Experimental timeline: Control or *Lphn2ECKO* pups were injected with 100µg of tamoxifen at postnatal day 1, 2 and 3 (P1, P2 and P3); BMP9/10 blocking antibodies or IgG control antibody at P4; and retinas were harvested at P7. **B**) Representative images of one quadrant of P7 retinas from pups injected with control or BMP9/BMP10-blocking antibodies. Yellow arrow indicates arteriovenous malformations (AVMs). Scale bar, 500 µm. **C**) Quantification of AVMs at the vascular center in retinas from mice injected with control or BMP9/BMP10-blocking antibodies (n = 10, 12, 6 and 8 retinas for control-IgG, control-BMP9/10 blocking antibody, Lphn2ECKO-IgG and Lphn2ECKO-BMP9/10 blocking antibody, respectively) from (**B**). Values are means ± S.E.M. Statistical significance was calculated using *Student*’s t-test. (**D**) Quantification of vascular branch points in Lphn2 ECKO P7 retinas. Branch points are normalized to area as in Figure 8F. Statistics used one-way ANOVA with post-hoc Tukey test.

**Supplementary figure 7:**
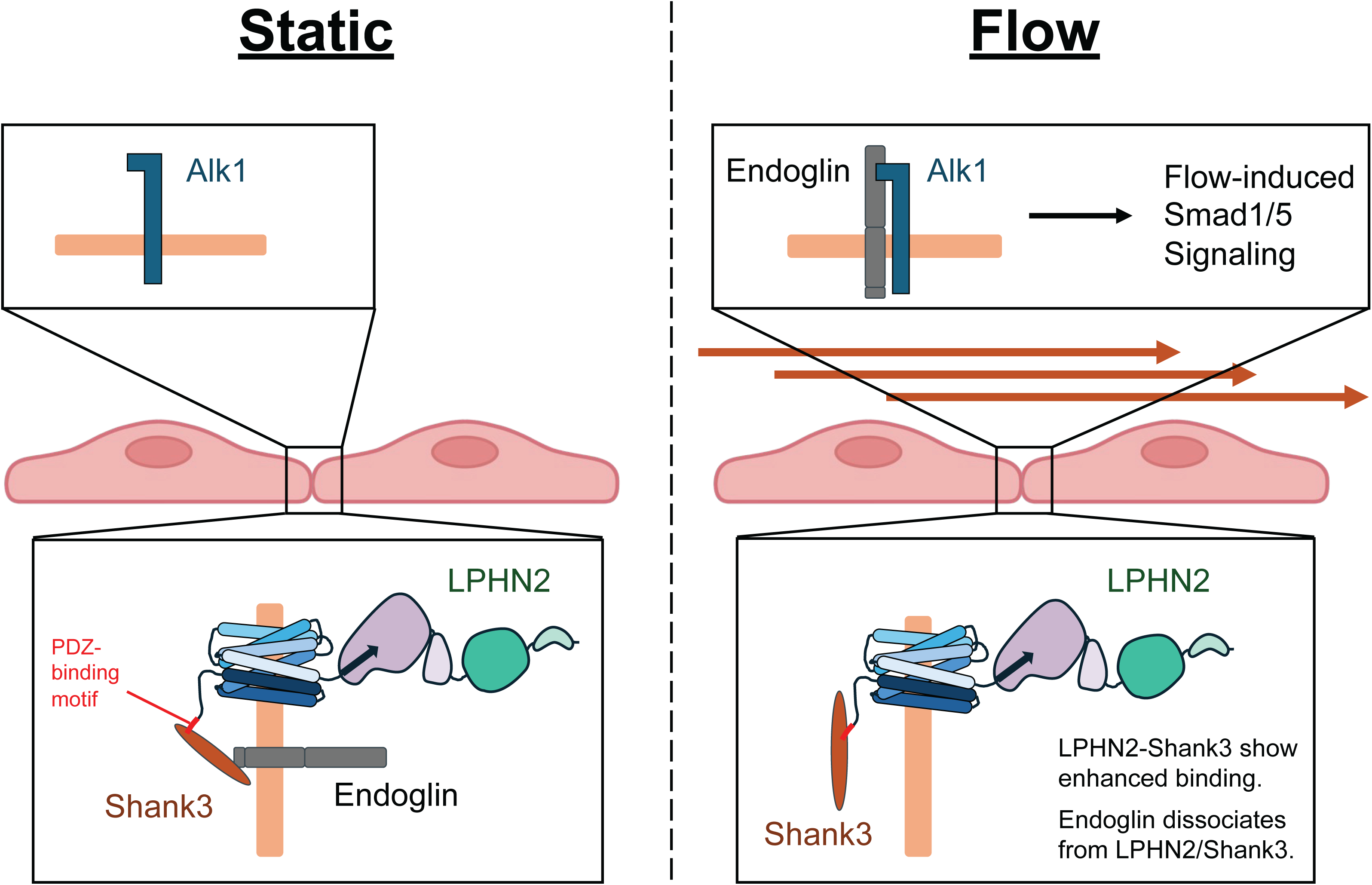
Schematic for LPHN2/Shank3/Endoglin dynamics and Smad1/5 activation Under static condition, LPHN2 and Shank3 sequester Endoglin. Upon the onset of flow, Endoglin dissociates from the LPHN2/Shank3 complex and associates with Alk1, triggering downstream Smad1/5 signaling.

## Acknowledgements

This work was supported by NIH grant R01 HL155543 to MAS and AE. We thank Prof. Yong-Hui Jiang and Dr. Sheng-Nan Qiao at Yale school of medicine for Shank3 global knockout mice. We thank Dr. Ross Tomaino at the Taplin Mass spectrometry facility, Harvard University.

## Author contributions

KT and MAS conceptualized the study. KT, MAS and MChanduri designed the experiments. KT and MChanduri carried out the experiments and analyzed the data. HP performed and analyzed the mouse AVM experiments. MChen and AP performed *in vivo* experiments in Lphn2 ECKO mice and zebrafish, respectively. DJ assisted injections in the AVM study. YMK performed an imaging-based *in vitro* experiment and validation of gene knockdowns. JS assisted *in vitro* experiment. AE provided financial support and with manuscript revisions. MChanduri and MAS wrote the first draft. KT, MChanduri and MAS edited the manuscript.

